# Cocaine, via ΔFosB, remodels gene expression and excitability in ventral hippocampus

**DOI:** 10.1101/2024.11.06.621895

**Authors:** Andrew L. Eagle, Marie A. Doyle, Chiho Sugimoto, Megan Dykstra, Hayley M. Kuhn, Brooklynn R. Murray, Ryan M. Bastle, He Jin, Ian Maze, Michelle S. Mazei-Robison, Alfred J. Robison

## Abstract

Ventral hippocampus (vHPC) CA1 pyramidal neurons send glutamatergic projections to nucleus accumbens (NAc), and this vHPC-NAc circuit mediates cocaine seeking and reward, but it is unclear whether vHPC-NAc neuron properties are modulated by cocaine exposure to drive subsequent behavior. The immediate early gene transcription factor ΔFosB is induced throughout the brain by cocaine and is critical for cocaine seeking, but it’s function in vHPC-NAc neurons is not understood. We now show that circuit-specific knockout of ΔFosB in vHPC-NAc neurons impaired cocaine reward expression and forced abstinence-induced seeking. We also found that vHPC-NAc excitability was decreased by experimenter-administered repeated cocaine and cocaine self-administration, and this cocaine-induced excitability decrease was mediated by ΔFosB expression. To uncover the mechanism of this change in circuit function, we used circuit-specific translating ribosome affinity purification (TRAP) to assess cocaine-induced, ΔFosB-dependent changes in gene expression in vHPC-NAc. We found that cocaine causes a ΔFosB-dependent increase in the expression of calreticulin, an ER-resident calcium-buffering protein. Calreticulin expression mediated vHPC-NAc excitability and was necessary for cocaine reward. These findings uncover a novel, non-canonical mechanism by which cocaine increases calreticulin in vHPC leading to decreased vHPC-NAc excitability and drives cocaine seeking and reward.

## Introduction

Addiction is a crucial social and health problem that can be defined as the unnatural drive to seek and take drugs despite negative consequences. Cocaine overdose deaths tripled between 2015 and 2020 (Statistics, 2021), yet there are currently no FDA-approved treatments for cocaine substance use disorders. To develop novel therapeutics for cocaine SUD, we require a better understanding of how drugs remodel neuronal circuitry that may underlie addictive behavior. Addiction is characterized by reward dysfunction and aberrant seeking behavior, e.g. relapse, leading to changes in neuroplasticity, gene expression, and neurocircuitry (Feltenstein, See, & Fuchs, 2020; Kalivas, 2009; Koob & Volkow, 2016; Lüscher & Malenka, 2011; Robison & Nestler, 2011; Volkow & Morales, 2015; Wolf, 2016). The ventral hippocampus (vHPC) is important for emotional memory and is functionally distinct from dorsal HPC (Fanselow & Dong, 2010; Strange, Witter, Lein, & Moser, 2014), and new evidence suggests a role for vHPC in encoding external stimuli with internal drives, including the motivation to seek reward (V. S. Turner, O’Sullivan, & Kheirbek, 2022). vHPC is necessary for the relapse to drug seeking (Bossert et al., 2016; Bossert & Stern, 2012; Fredriksson et al., 2021; Lasseter, Xie, Ramirez, & Fuchs, 2010; Marchant et al., 2016; Rogers & See, 2007; W. Sun & Rebec, 2003), and acute activation of vHPC induces drug seeking (Taepavarapruk, Butts, & Phillips, 2015; Taepavarapruk & Phillips, 2003; Vorel, Liu, Hayes, Spector, & Gardner, 2001). Glutamatergic afferents that innervate nucleus accumbens (NAc) medium spiny neurons (MSNs) are important in drug-induced plasticity, reward, and seeking (Britt et al., 2012; Kalivas, 2009; Stuber, Britt, & Bonci, 2012; Wolf, 2010, 2016; Zinsmaier, Dong, & Huang, 2022). vHPC sends glutamatergic projections to NAc (vHPC-NAc), and this circuit is critical for cocaine reward and seeking (Britt et al., 2012; LeGates et al., 2018; Sjulson, Peyrache, Cumpelik, Cassataro, & Buzsáki, 2018; V. S. Turner et al., 2022; Zhou et al., 2019). Through excitatory synaptic plasticity, cocaine promotes indelible changes that underlie addictive behavior, like altered reward and drug seeking. Indeed, cocaine reshapes the plasticity of vHPC inputs onto NAc MSNs (Britt et al., 2012; LeGates et al., 2018; Pascoli et al., 2014; Sjulson et al., 2018), and reversal of this plasticity impairs cocaine seeking (Pascoli et al., 2014). However, mechanisms by which cocaine remodels the function of vHPC neurons that project onto NAc MSNs remain unknown.

Psychostimulants, like cocaine and amphetamine, alter the intrinsic excitability of neurons in regions including ventral subiculum (Cooper, Moore, Staff, & Spruston, 2003), cortex (Nasif, Sidiropoulou, Hu, & White, 2005), and NAc MSNs (Delint-Ramirez, Garcia-Oscos, Segev, & Kourrich, 2020; Kalivas & Hu, 2006). Of note, amphetamine decreases the intrinsic excitability of neurons in ventral subiculum (Cooper et al., 2003), a region close in proximity to the vHPC that also sends projections to the NAc. Collectively, these findings suggest that not only do drugs of abuse produce changes in synaptic plasticity in accumbal synapses, but they also produce changes in brain regions projecting to NAc that regulate addictive behavior, e.g. vHPC. However, it is currently unknown whether cocaine directly affects the intrinsic extricability of vHPC-NAc cells.

Cocaine may be remodeling vHPC neurons via transcriptional changes. ΔFosB, a stable immediate early gene transcription factor (Robison & Nestler, 2011; Teague & Nestler, 2022) that mediates hippocampal neuron function (Eagle et al., 2015; Eagle et al., 2020; Eagle, Williams, Beatty, Cox, & Robison, 2018), is induced in vHPC by cocaine (Gajewski et al., 2019; Perrotti et al., 2008), and its expression in vHPC is necessary for cocaine reward (Gajewski et al., 2019). Thus, we hypothesized that cocaine, via ΔFosB, may be remodeling vHPC neurons through altered gene expression and cell excitability to drive cocaine reward and seeking behavior. Indeed, the current study demonstrates that cocaine induces ΔFosB, which drives expression of the target gene calreticulin to decrease vHPC-NAc excitability and drive cocaine reward and seeking.

## Results

### Cocaine increases ΔFosB expression in vHPC-NAc neurons

We have previously established that repeated, experimenter-administered cocaine induces ΔFosB mRNA and protein expression in vHPC neurons (Gajewski et al., 2019). There is also semi-quantitative evidence suggesting ΔFosB is induced by cocaine self-administration in dorsal HPC (Perrotti et al., 2008). We sought to determine the extent to which cocaine self-administration induces ΔFosB in vHPC neurons. Separate cohorts of male mice were trained to self-administer IV infusions of cocaine (0.5 mg/kg per infusion) or saline for 14 d (Fig 1a-b; Suppl. Fig. 1a-b). One day after the last session, vHPC tissue was taken for assessment of ΔFosB mRNA and protein. Cocaine self-administration increased ΔFosB mRNA in vHPC (Fig. 1c) compared to saline self-administering mice.

**Figure 1.**
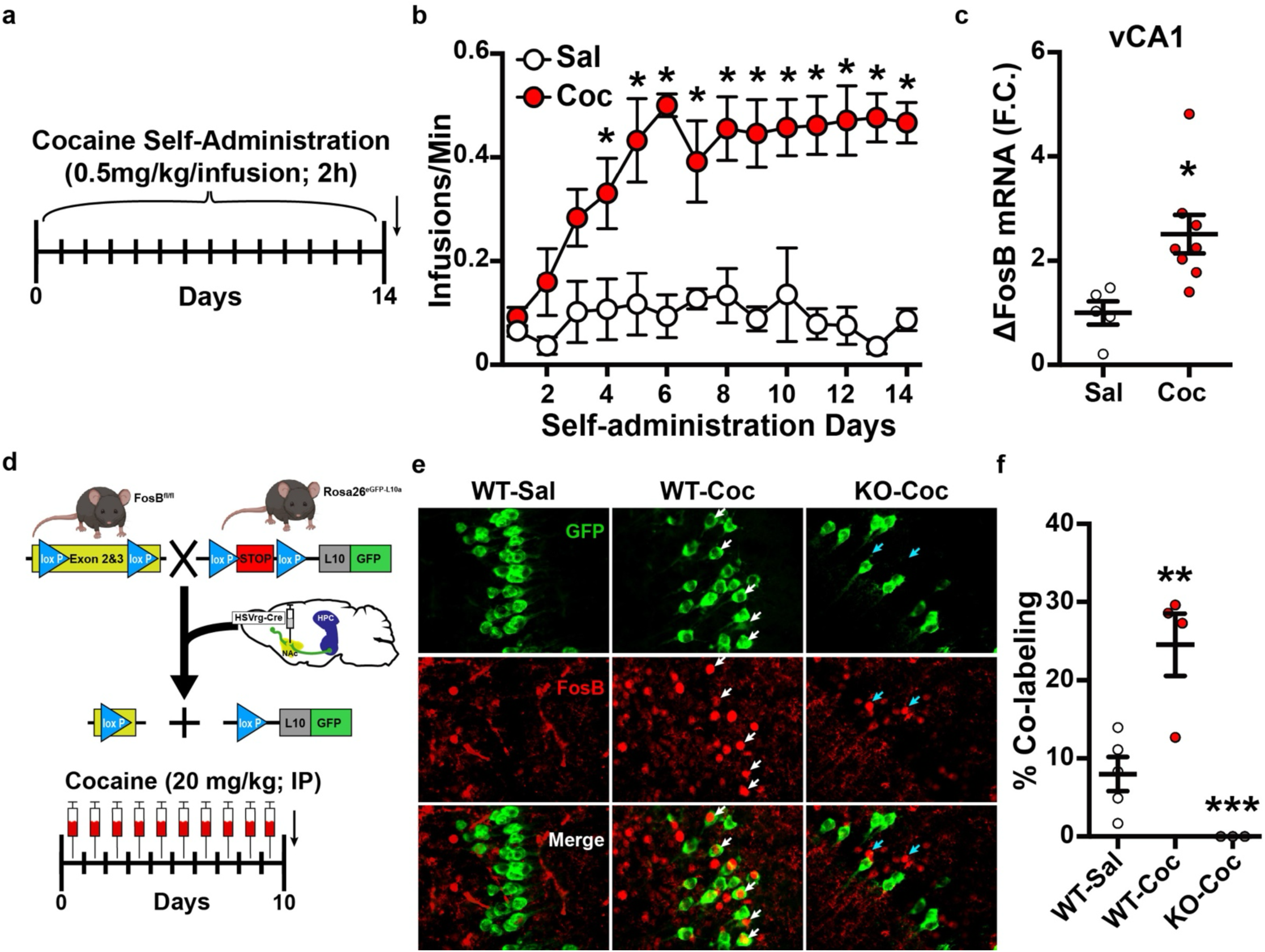
Chronic cocaine induces ΔFosB in ventral hippocampus neurons that project to nucleus accumbens. **a)** Timeline of cocaine or saline self-administration in male mice. Arrow indicates the time of tissue collection. **b)** Cocaine self-administration performance. Cocaine or saline infusions per minute across 14 daily 2 h sessions where responses on an active nosepoke resulted in an IV infusion of cocaine (Coc; 0.5 mg/kg/infusion; n=8) or saline (Sal; n=5). Cocaine self-administering mice received greater infusions across days compared to saline self-administering mice [Two-way RM ANOVA: DayXGroup F (13, 143) = 2.911, P=0.009], with significantly more infusions on days 4-14 (Holm-Sidak tests *P<0.05 for days 4-14). **c)** Cocaine self-administration enhances ΔFosB in vHPC. vHPC tissue was taken 1 d after the last self-administration session for detection of ΔFosB mRNA or protein. Cocaine self-administration significantly increased ΔFosB mRNA compared to saline [Independent samples t-test: t(11)=2.997, *P=0.0121]. **d)** Circuit-tagging procedure. Male mice with crossed Floxed FosB and L10-GFP (KO; *FosB^fl/fl^*:*Rosa26^eGFP-L10a^*) or L10-GFP only (WT; *Rosa26^eGFP-L10a^*) received retrograde HSV vector expressing Cre (HSVrg-hEf1ɑ-Cre) into nucleus accumbens to drive GFP and FosB KO in projections to accumbens, including vHPC-NAc projections. All mice were treated with cocaine (Coc; 20 mg/kg; IP) or saline (Sal) for 10 d and 1 d later vHPC coronal brain slices were taken for FosB/ΔFosB immunohistochemistry. **e**) Representative 20x coronal images of GFP (green, top), FosB/ΔFosB (red, bottom), and a merged image (bottom) in WT-Sal (n=5), WT-Coc (n=4), and KO-Coc (n=3) mice. White arrows denote GFP+ neurons co-expressing FosB/ΔFosB. **f)** Cocaine increases ΔFosB in vHPC-NAc neurons. Quantification of co-labeled GFP+ and FosB/ΔFosB+ neurons in all three groups. WT-Coc had significantly greater co-labeled neurons compared to WT-Sal and KO-Coc [One-way ANOVA: F (2, 9) = 18.01, P=0.0007; Holm-Sidak **P<0.01, ***P<0.001].

The vHPC contains heterogenous cell types, but NAc-projecting vHPC neurons (vHPC-NAc), which are primarily in vCA1 and ventral subiculum (vSub; Eagle et al., 2020), are critical for cocaine seeking (Britt et al., 2012; Pascoli et al., 2014). We assessed whether repeated cocaine increases ΔFosB expression in vHPC-NAc neurons using a circuit-tagging approach (Fig. 1d) to drive the expression of GFP and knockout the *FosB* gene in projections to NAc. Briefly, Cre-dependent GFP reporter mice (*Rosa26^eGFP-L10a^*) were crossed with floxed *FosB* mice (*FosB^fl/fl^:Rosa26^eGFP-L10a^*). Both wild-type (WT; *Rosa26^eGFP-L10a^*) and conditional knockout mice (FosB KO; *FosB^fl/fl^:Rosa26^eGFP-L10a^*) received intracranial infusions of a retrograde HSV vector expressing Cre recombinase (HSVrg-hEf1α-Cre) into NAc. Three to six weeks later, circuit-tagged WT and FosB KO mice received 10 d of repeated experimenter-administered cocaine (20 mg/kg; IP) or saline and were assessed for FosB/ΔFosB immunohistochemistry in GFP-tagged vHPC-NAc neurons. Similar to our previous findings in whole vHPC (Gajewski et al., 2019), cocaine increased ΔFosB expression in vHPC-NAc neurons (Fig. 1e-f). This effect was completely blocked in vHPC-NAc neurons of the FosB KO mice. These data show that experimenter- or self-administered cocaine increase ΔFosB in vHPC, and specifically that cocaine induces ΔFosB in vHPC-NAc neurons that are important for drug seeking.

### ΔFosB in vHPC-NAc is necessary for cocaine reward and seeking

vHPC neurons, including those that project to NAc, are critically important for drug seeking (Bossert et al., 2016; Bossert & Stern, 2012; Britt et al., 2012; Fredriksson et al., 2021; Lasseter et al., 2010; Marchant et al., 2016; Pascoli et al., 2014; Rogers & See, 2007; W. Sun & Rebec, 2003; Vorel et al., 2001) and reward (LeGates et al., 2018; Sjulson et al., 2018; Zhou et al., 2019). Based on our finding that cocaine induces ΔFosB in vHPC-NAc neurons, we surmised that ΔFosB may be playing a role in vHPC-NAc drive of cocaine seeking and reward. We have previously demonstrated that hippocampal ΔFosB expression regulates learning and memory (Eagle et al., 2015) and cocaine CPP (Gajewski et al., 2019), but it’s role in vHPC-NAc neurons in the context of drug seeking remains unknown. Therefore, we next sought to determine whether ΔFosB expression in vHPC-NAc is necessary for cocaine reward using our previously characterized circuit-specific CRISPR approach (Fig. 2a) that knocks down ΔFosB specifically in vHPC-NAc neurons (Eagle et al., 2020). CRISPR-mediated FosB KO (FosB crKO) significantly reduced the preference for a context conditionally paired with cocaine (Fig. 2b-d), demonstrating that ΔFosB expression in vHPC-NAc neurons is necessary for the expression of cocaine reward. However, place preference does not distinguish between deficits in the processing of reward or deficits in ability to form context-reward association memories. To determine the specific role of vHPC-NAc ΔFosB expression, we turned to an operant cocaine self-administration model.

**Figure 2.**
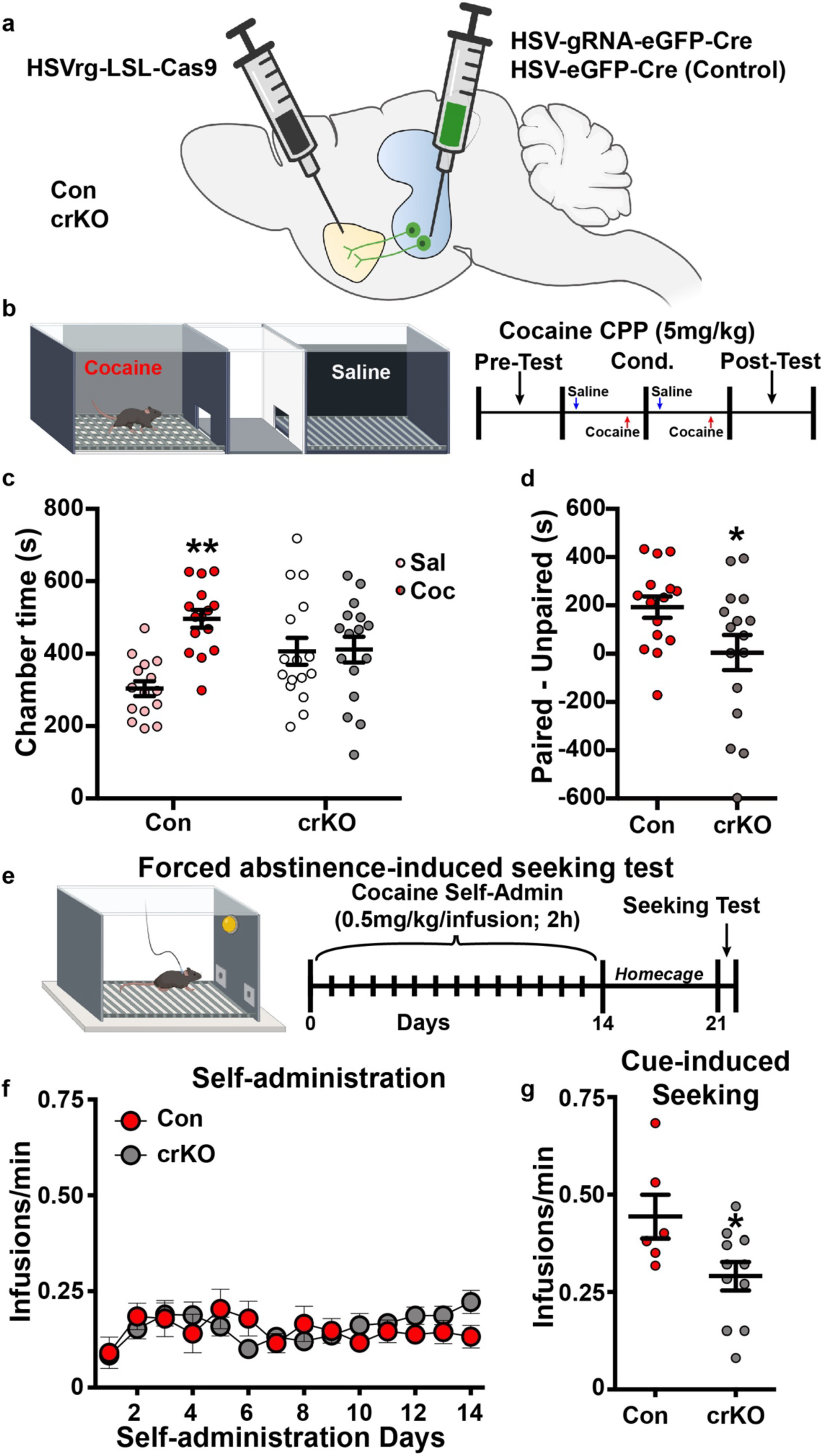
Selective FosB knockout in ventral hippocampus neurons that project to nucleus accumbens impairs cocaine reward and cocaine seeking. **a)** Illustration of dual virus CRISPR method to selectively knockout FosB/ΔFosB in vHPC-NAc neurons. **b)** Cocaine CPP procedure and timeline. Male controls (Con; n=15) or FosB crKO (crKO; n=16) mice underwent cocaine CPP in a 3-chamber box for twice-daily conditioning sessions of saline and cocaine (5 mg/kg; IP) followed by a drug-free post-test. **c)** Saline-paired (Sal) and cocaine-paired (Coc) chamber time during the post-test for Con and crKO mice. Con mice spent significantly more time in the Coc chamber compared to the Sal chamber, however crKO had no differences in time spent between chambers [Two-way RM ANOVA: ChamberXGroup: F (1, 29) = 4.738, P=0.0378; Holm-Sidak **P<0.01]. **d)** Cocaine-paired and cocaine-unpaired (i.e., saline) chamber difference in Con and crKO mice. crKO had significantly reduced paired-unpaired time compared to Con [Independent samples t-test: t(29)=2.177, *P=0.0378]. **e)** Forced abstinence-induced seeking procedure. **f)** Con (n=6) and crKO (n=12) mice did not differ in the number of infusions across cocaine self-administration sessions. **g)** After a 14 d forced abstinence in their homecage, Con increased the number of mock infusions per minute during a seeking test compared to crKO [Independent samples t-test: t(15)=2.377, *P=0.0312].

Forced abstinence from cocaine self-administration produces long-term changes in plasticity (Ma et al., 2014), including at vHPC synapses onto NAc MSNs (Pascoli et al., 2014), and reversing these changes attenuates cocaine seeking (Pascoli et al., 2014; Pascoli, Turiault, & Lüscher, 2012). Therefore, we next asked whether ΔFosB expression in vHPC-NAc is necessary for acquisition and maintenance of cocaine self-administration and/or for forced abstinence-induced cocaine seeking. Mice were trained to respond on an active nosepoke for cocaine for 14 d (0.5 mg/kg/infusion; IV) on a fixed ratio FR1 schedule (Fig. 2e). FosB crKO had no effect on the acquisition or maintenance of cocaine self-administration, indicated by number of infusions (Fig. 2f), nosepoke responses, and accuracy (Suppl. Fig. 2a,c). Mice were then left in their homecages for 1 week before testing for abstinence-induced seeking back in the cocaine context. In the seeking test, nosepoke responses only resulted in mock infusions of saline and the sound of the infusion pump. We found that vHPC-NAc FosB crKO impaired abstinence-induced cocaine seeking, without affecting cocaine self-administration (Fig. 2g; Suppl. Fig. 2b,d), reducing the number of mock infusions (Fig. 2g) and active nosepoke responses (Suppl. Fig. 2b), without affecting accuracy (Suppl. Fig. 2d). Collectively, these findings indicate that ΔFosB expression in vHPC-NAc is required for context-induced cocaine reward and seeking.

### ΔFosB mediates cocaine-induced decreases in vHPC-NAc excitability

Psychostimulants, like cocaine and amphetamine, alter the intrinsic excitability of neurons in ventral subiculum (Cooper et al., 2003), cortex (Nasif et al., 2005; Otis et al., 2018), lateral habenula (Neumann et al., 2015), and NAc (Delint-Ramirez et al., 2020; Dong et al., 2006; Kalivas & Hu, 2006; Ma et al., 2013; Mu et al., 2010). Of particular importance, amphetamine decreases the intrinsic excitability of neurons in ventral subiculum (Cooper et al., 2003), a region close in proximity to the vHPC that also sends projections to the NAc (Boxer, Kim, Dunn, & Aoto, 2023). Collectively, these findings suggest that, beyond changes in synaptic plasticity at accumbal synapses, psychostimulants produce intrinsic excitability changes in brain regions that regulate drug responses and project to NAc. However, it is unknown whether cocaine directly affects the intrinsic extricability of vHPC neurons. To address this, male C57Bl/6J mice were trained to self-administer cocaine (0.5 mg/kg/infusion) or saline for 14 d on an FR1 active nosepoke schedule of reinforcement (Suppl. Fig. 3). One day after the last cocaine (or saline) self-administration session, vHPC was sliced for *ex vivo* electrophysiology, and we found that cocaine self-administration reduced vHPC CA1 pyramidal neuron excitability (Fig. 3a-d; Suppl. Table 1). This was indicated by a decrease in depolarization-induced spike frequency (Fig. 3b), a decrease in total spikes (Fig. 3c), and no changes in rheobase current (Fig. 3d). We also used circuit-tagging in cocaine and saline self-administering WT male mice (Fig. 3e) to assess excitability of ventral CA1 neurons that project to NAc (vCA1-NAc). vHPC tissue was taken 1 d after the last self-administration session and we found that cocaine self-administration also reduced vCA1-NAc excitability (spike frequency and total spikes) (Fig. 3f-g; Suppl. Table 2). We also observed an increase in rheobase in vCA1-NAc neurons from cocaine self-administering mice (Fig. 3h). We previously established that ΔFosB expression decreases dorsal and ventral CA1 pyramidal neuron excitability (Eagle et al., 2020; Eagle et al., 2018), and ΔFosB knockout enhances vCA1-NAc excitability in male mice (Eagle et al., 2020). Based on our finding that cocaine induces ΔFosB in vHPC-NAc neurons, we hypothesized that ΔFosB expression mediates the cocaine-dependent reduction in vCA1-NAc excitability. We used our circuit tagging approach (Fig. 3i) to express GFP and knock out ΔFosB in vCA1-NAc neurons in Cre-dependent GFP (WT) and floxed FosB (FosB KO) male mice treated with 10 d of experimenter-administered cocaine (20 mg/kg; IP) or saline. vHPC slices were taken 1 d after the last injection for whole-cell electrophysiology. We found that experimenter-administered cocaine in WT mice reduced vCA1-NAc excitability, as indicated by a decrease in depolarization-induced spike frequency (Fig. 3j-k), with no change in rheobase (Fig. 3l), or other intrinsic cellular properties (Suppl. Table 3). Critically, ΔFosB knockout in vCA1-Nac neurons (FosB KO mice) blocked this cocaine-induced decrease in excitability. These findings collectively suggest that cocaine reduces the excitability of vHPC-NAc neurons via induction of ΔFosB.

**Figure 3.**
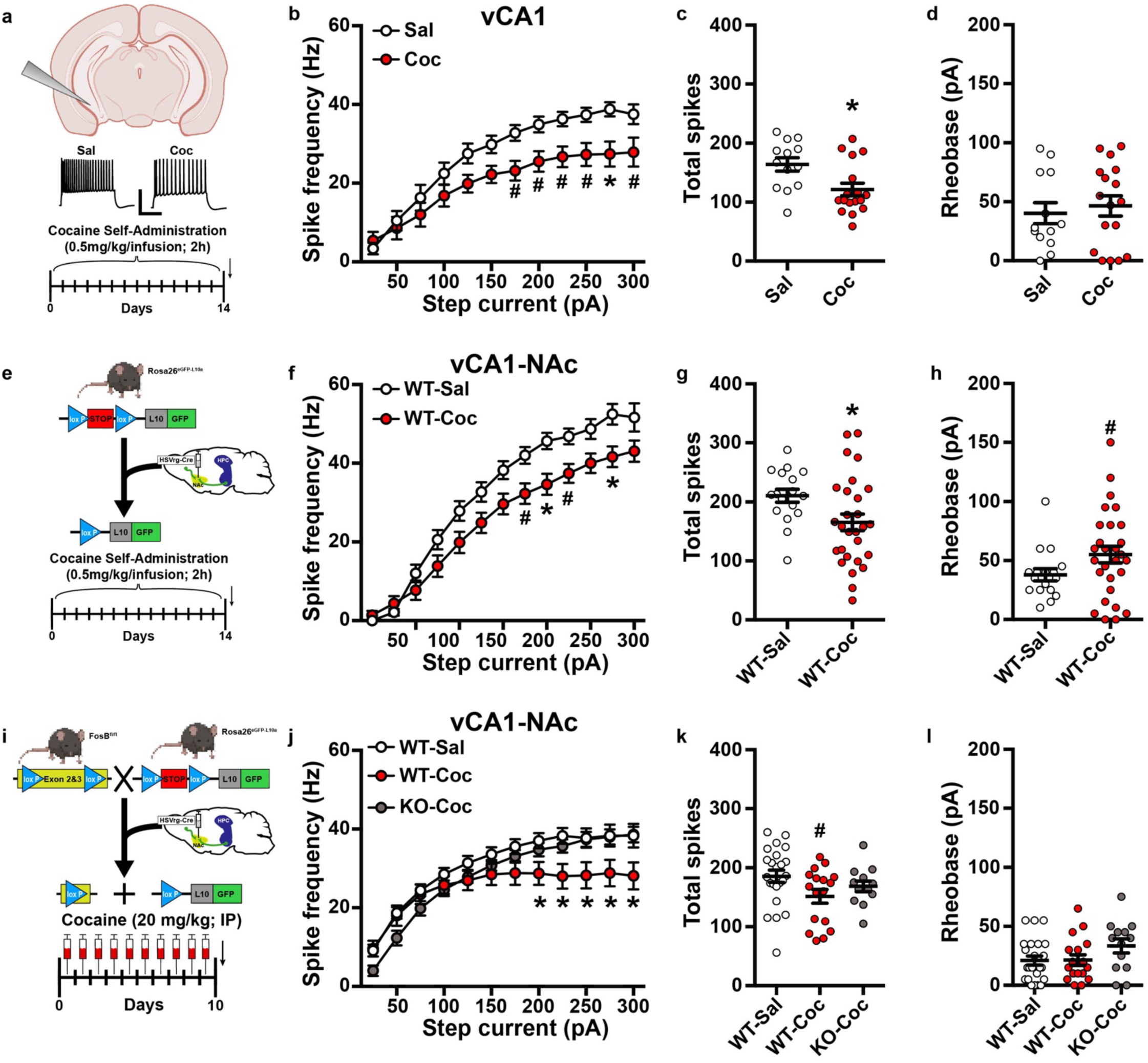
Cocaine, through ΔFosB, decreases the membrane excitability of ventral hippocampus neurons that project to nucleus accumbens. **a)** Illustration of vHPC *ex vivo* patch-clamp approach (top) and representative spike traces at 200 pA in vCA1 neurons of cocaine (Coc; n=4 mice; n=17 cells) and saline (Sal; n=3; n=13 cells) self-administering male mice. Scale bar 50pA, 200 ms. **b)** Cocaine self-administration reduces vCA1 neuron spike frequency across depolarizing steps of current [Two-way RM ANOVA: ME of Group F (1, 28) = 7.320, P=0.0115; CurrentXGroup F (11, 308) = 1.993, P=0.0286; Holm-Sidak *P<0.05, ^#^P<0.10]. **c)** Cocaine self-administration reduces the total number of spikes [Independent samples t-test: t(28)=2.705, *P=0.0115]. **d)** There were no significant differences between Coc or Sal groups on rheobase. **e)** Circuit-tagging procedure (top; from Fig. 1f) and experimenter-administered cocaine timeline in male mice (bottom). WT mice received either saline (WT-Sal; n=8 mice; n=23 cells) or cocaine (WT-Coc; n=6 mice; n=17 cells). FosB KO mice received cocaine (KO-Coc; n=4 mice; n=13 cells). **f)** Cocaine significantly reduced spike frequency across depolarizing steps of current in WT mice [Two-way RM ANOVA: CurrentXGroup F (22, 550) = 3.545, P<0.0001], specifically at current steps of 200-300pA [Holm-Sidak WT-Sal vs WT-Coc *P<0.05 for 200-300 pA]. There were no differences between WT-Sal and KO-Coc. KO-Coc had reduced spike frequency across steps compared to WT-Coc at current steps of 225-300 pA [Holm-Sidak WT-Coc vs KO-Coc **^‡^**P<0.05 for 225-300 pA]. **g)** There was a trend towards a significant difference in the total number of spikes in WT-Coc vCA1-NAc neurons compared to WT-Sal and KO-Coc neurons [One-way ANOVA: F (2, 50) = 2.728, P=0.0751; Holm-Sidak ^#^P<010]. **h)** There were no differences between groups on rheobase. **i)** Circuit-tagging procedure (top; from Fig. 1f) and cocaine (WT-Coc; n=6 mice; n=28 cells) or saline (WT-Sal; n=5 mice; n=17 cells) self-administration timeline in male mice (bottom). **j)** Cocaine self-administration significantly reduced spike frequency across depolarizing steps of current in vCA1-NAc neurons [Two-way RM ANOVA: CurrentXGroup F (12, 516) = 4.674, P<0.0001; Holm-Sidak *P<0.05, ^#^P<010]. **k)** Cocaine self-administration significantly reduced the total number of spikes compared to saline self-administration [Independent samples t-test: t(43)=2.230, *P=0.0310]. **l)** There was a slight trend towards a significant increase in rheobase in vCA1-NAc neurons of cocaine self-administering mice [Independent samples t-test: t(43)=1.719, ^#^P=0.0928].

### Chronic excitability underlies cocaine reward

Cocaine-induced changes in plasticity underlie many behavioral responses to the drug (Dong et al., 2006; Ma et al., 2014; Pascoli et al., 2014; Pascoli et al., 2012). Therefore, we hypothesized that cocaine-induced reduction in vHPC-NAc excitability underlies cocaine reward and seeking behaviors. Acute activation of vHPC, including vHPC-NAc neurons, enhances or induces cocaine seeking (Vorel et al., 2001), whereas acute inhibition can decrease cocaine seeking (Britt et al., 2012). However, it is unclear how a chronic decrease in membrane excitability underlies cocaine behavioral responses. Reversal of drug-induced changes in prefrontal cortex and nucleus accumbens neuronal excitability ameliorates cocaine seeking (B. T. Chen et al., 2013; Dong et al., 2006; Otis et al., 2018; Yousuf et al., 2019). In line with this evidence, we assessed whether chronic chemogenetic increase in excitability of vHPC-NAc neurons affects cocaine reward (Fig. 4a-b). To express circuit-specific DREADD Gq receptors specifically in vHPC-NAc neurons, we used an intersectional virus strategy (Fig. 4a). Male C57Bl/6J mice received infusions of retrograde HSV expressing Cre into NAc (HSVrg-hEf1α-Cre) and either AAV2-CMV-GFP or AAV2-hSyn-DIO-hM3Dq-mCherry in vHPC. To chronically activate vHPC-NAc neurons, mice received 14 d of twice-daily i.p. injections of the DREADD ligand, CNO (0.3 mg/kg). During the last 4 days, mice underwent cocaine CPP (Fig. 4b). Chronic vHPC-NAc activation impaired place preference for cocaine (Fig. 4c-d). Specifically, GFP control mice spent significantly more time in a chamber paired with cocaine compared to a chamber paired with saline (Fig. 4c). However, Gq-expressing mice spent a similar amount of time in both the saline- and cocaine-paired chambers. Preference (cocaine-paired minus cocaine-unpaired chamber) was also significantly reduced in Gq mice compared to GFP mice (Fig. 4d). Interestingly, a shorter subchronic activation (4 d of CNO treatment during the CPP) did not impair the expression of cocaine reward (Suppl. Fig. 4). These findings demonstrate that cocaine-induced decreases in vHPC-NAc excitability are necessary for cocaine place preference.

**Figure 4.**
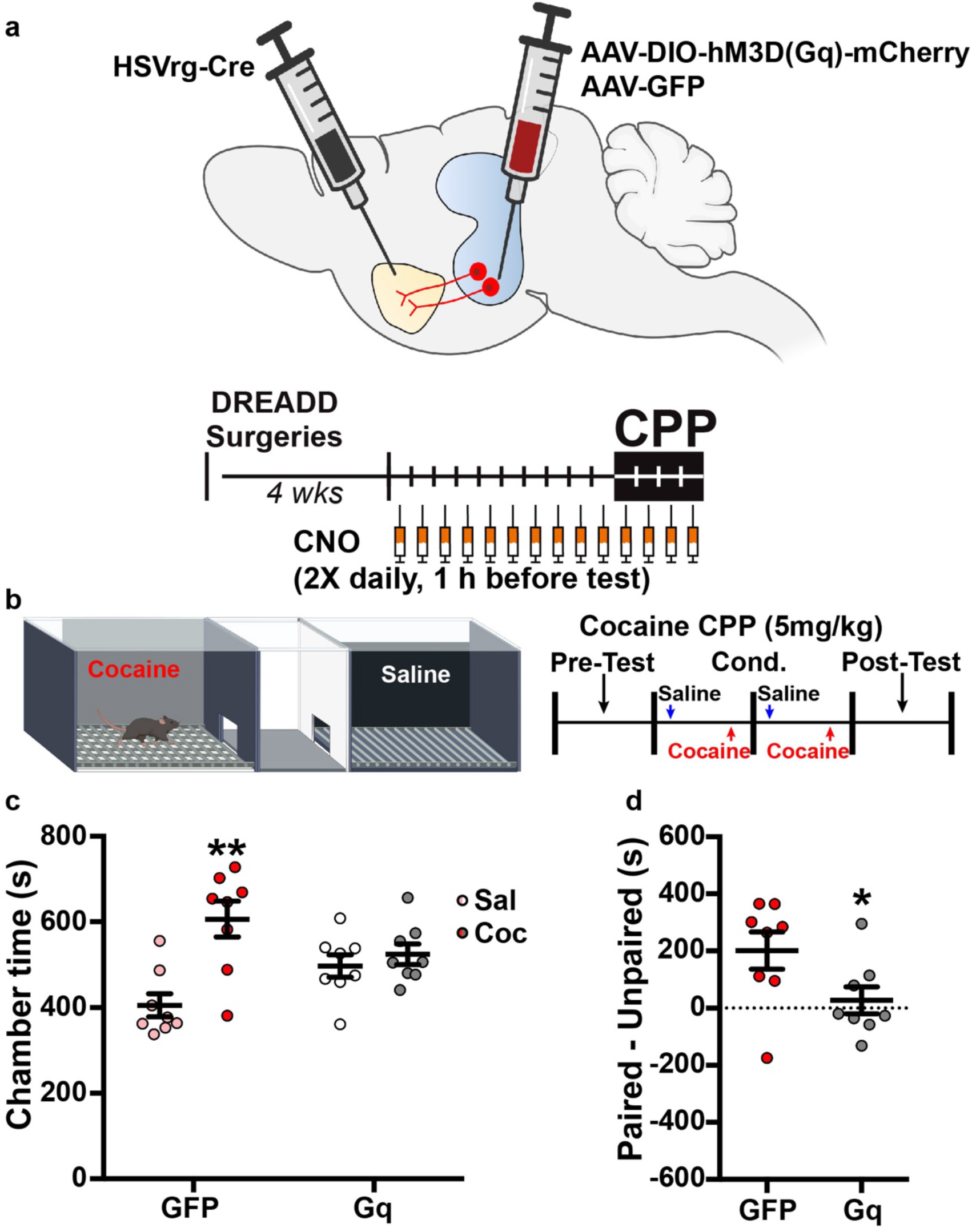
Chronic chemogenetic activation of ventral hippocampus neurons that project to nucleus accumbens impairs the conditioned place preference for cocaine. **a)** Viral approach to express Gq receptors in vHPC-NAc neurons of male mice. A retrograde HSV vector expressing Cre (HSVrg-hEf1ɑ-Cre) was injected into nucleus accumbens and either a control (AAV2-CMV-GFP) or DREADD Gq viral vector (AAV2-hSyn-DIO-hM3Dq-mCherry) was injected into ventral hippocampus. **b)** Cocaine CPP procedure and timeline. CNO was administered twice-daily for 14 d that ended with cocaine (5 mg/kg; IP) CPP test. **c)** Saline-paired (Sal) and cocaine-paired (Coc) chamber time during the post-test for GFP (n=8) and Gq (n=8) mice. GFP mice spent significantly more time in the Coc chamber compared to the Sal chamber, however Gq did not differ in time spent between chambers [Two-way RM ANOVA: ChamberXGroup: F (1, 14) = 4.737, P=0.0471; Holm-Sidak **P=0.0063]. **d)** Cocaine-paired and cocaine-unpaired (i.e., saline) chamber difference in GFP and Gq mice. Gq had significantly reduced paired-unpaired time compared to GFP [Independent samples t-test: t(14)=2.176, *P=0.0471].

### Cocaine, via ΔFosB, alters gene expression in vHPC-NAc neurons

ΔFosB regulates gene transcription in a variety of brain regions, including the hippocampus (Eagle et al., 2020; Nestler, Kelz, & Chen, 1999; Robison & Nestler, 2011; Teague & Nestler, 2022), therefore we next sought to identify a transcriptional mechanism of cocaine-induced changes in membrane excitability and cocaine behavior. We employed a transcriptomic approach in vHPC-NAc using TRAP-Seq (Translating Ribosomal Affinity Purification followed by sequencing) (Allison et al., 2015) to assess actively translating mRNAs in vHPC-NAc neurons (Fig 5a). Circuit-tagged WT and KO male mice were experimenter-administered 10 d of cocaine (20 mg/kg; IP) or saline. One day after the last injection, fresh frozen vHPC tissue punches containing GFP+ cells were collected, samples were pooled (n=4-5/sample) and processed for TRAP-Seq. We first compared WT mice that received cocaine or saline. We plotted all 14,495 genes across confidence (P-value) and fold change (Fig. 5b). We identified 587 total significant cocaine-regulated genes (*p*<0.05), 270 genes that were upregulated by cocaine and 317 that were downregulated by cocaine. We also compared mice treated with cocaine with intact FosB (WT) and FosB KO in vHPC-NAc neurons (Fig. 5c). We identified 774 total significant cocaine-regulated genes (*p*<0.05), 385 genes that were upregulated by cocaine and 390 that were downregulated by cocaine. Of these significantly regulated genes we examined cocaine-regulated ΔFosB-dependent genes. We defined these genes as those that were reversibly regulated; either genes upregulated by cocaine in WT mice, and downregulated by FosB KO in cocaine-treated mice, or vice versa. We identified a total of 152 reversibly regulated genes. Of these, 108 were downregulated by cocaine and upregulated when FosB was knocked out in cocaine-treated mice, and 45 were upregulated by cocaine and downregulated when FosB was knocked out in cocaine-treated mice. These data revealed cocaine-induced, ΔFosB-dependent changes in actively translating gene expression.

**Figure 5.**
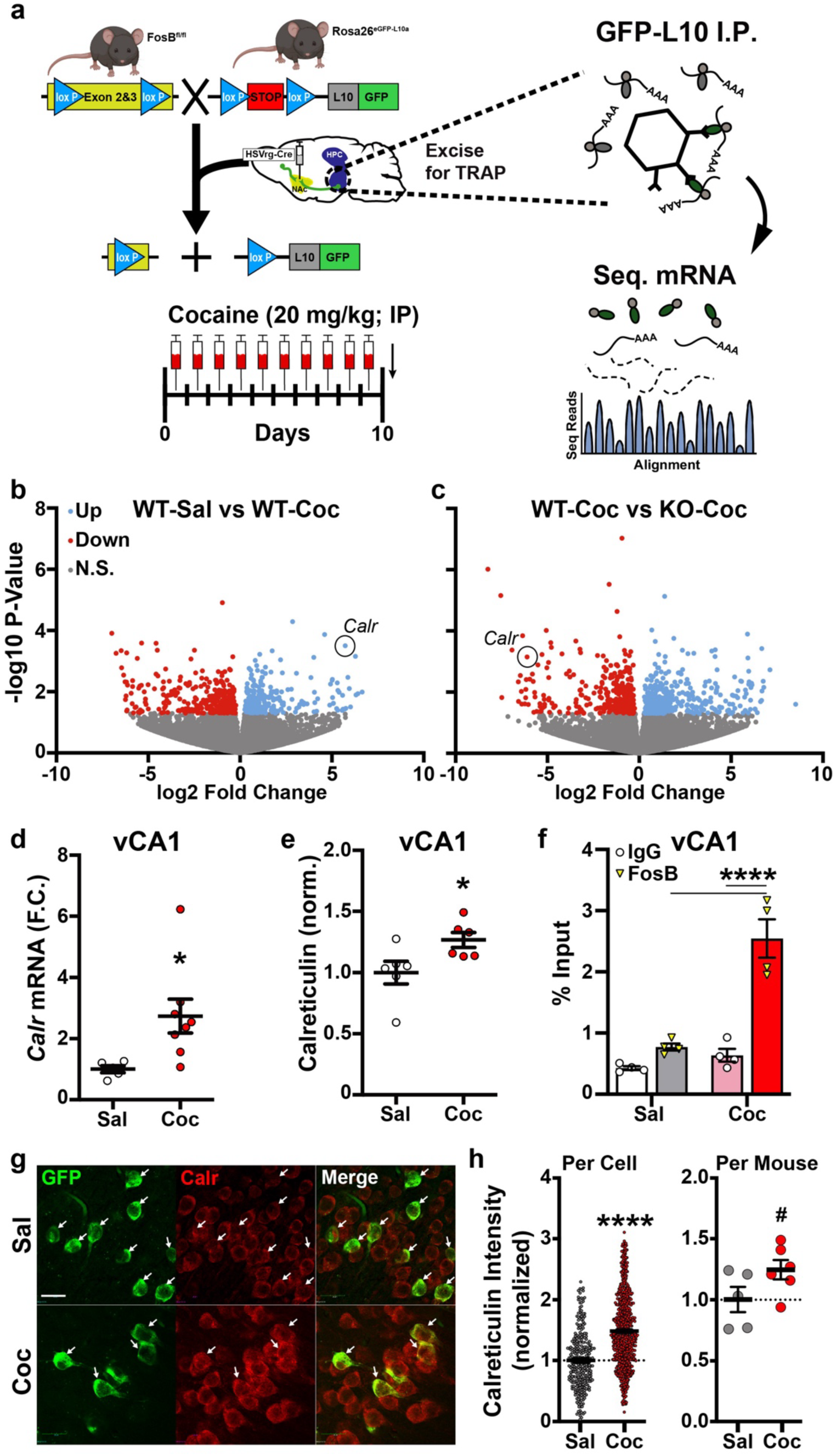
Cocaine-induced gene expression in ventral hippocampus neurons that project to nucleus accumbens. **a)** Circuit-tagging and TRAP-Seq approach (modified from(Eagle et al., 2020)). Circuit-tagged WT and FosB KO (KO) male mice received 10 d of cocaine (Coc; 20 mg/kg; IP) or saline (Sal). 1 d after the last injection, tissue punch samples containing GFP-tagged neurons were collected from WT-Sal (n=19), WT-Coc (n=19), and KO-Coc (n=20) mice. Samples were pooled 4-5 mice per sample (n=3-4 samples per group) and processed for TRAP-Seq. **b)** Volcano plot of differentially regulated genes in vHPC-NAc neurons between WT-Coc vs. WT-Sal mice. Gray indicates non-significant (N.S.) genes. **c)** Volcano plot of differentially regulated genes in vHPC-NAc neurons between KO-Coc vs. WT-Coc mice. Gray indicates non-significant (N.S.; P>0.05) genes. **d)** vHPC tissue punches from cocaine (Coc; n=8) and saline (Sal; n=5) self-administering male mice were processed for PCR for *Calr*. Cocaine self-administration increases *Calr* mRNA in vHPC [Independent samples t-test: t(11)=2.424, *P=0.0338]. **e)** vHPC tissue punches from cocaine (Coc, C; n=6) and saline (Sal, S; n=6) self-administering male mice were processed for Western blotting for calreticulin. Cocaine self-administration increases calreticulin in vHPC [Independent samples t-test: t(10)=2.428, *P=0.0356]. **f)** Male mice received 10 d of cocaine (Coc; 20 mg/kg; IP; n=12) or saline (Sal; n=12). Pooled vHPC tissue samples (n=3 mice/sample; n=4 samples per group) were processed for FosB/ΔFosB ChIP and PCR for *Calr*. IgG was used as control. Cocaine increased the binding of FosB/ΔFosB (% input) to the *Calr* promoter [Two-way ANOVA: InputXGroup: F (1, 12) = 21.64, P=0.0471; Holm-Sidak ****P<0.0001]. **g)** Circuit tagged male mice received 10 d of cocaine (Coc; 20 mg/kg; IP; n=6) or saline (Sal; n=5). 1 d after the last injection, perfused vHPC slices were collected for immunohistochemistry. Representative 40x images of calreticulin (red), GFP (green), and merged from coronal slices of vHPC. White arrows denote GFP+ vHPC-NAc neurons. **h)** Cocaine increased calreticulin intensity in GFP+ neurons [Independent samples t-test: t(1038)=12.75, ****P<0.0001]. **i)** There was a trend towards cocaine-induced increases in calfreticulin intensity in GFP+ neurons averaged per mouse [Independent samples t-test: t(9)=1.911, ^#^P=0.0883].

Of particular interest in these potential ΔFosB target genes was the ER-resident chaperone protein, calreticulin. Calreticulin is a high capacity, low affinity Ca^2+^-binding chaperone protein that, along with calnexin, regulates ER Ca^2+^ storage (Marek Michalak, Groenendyk, Szabo, Gold, & Opas, 2009). Calreticulin was upregulated in vHPC-NAc neurons by cocaine, but was downregulated in cocaine-treated FosB KO mice compared to WT. Conversely, when we knockout FosB in these same neurons, cocaine no longer increased calreticulin expression. This suggests that calreticulin expression may be increased by cocaine-induced ΔFosB. To verify our sequencing results, we first assessed whether cocaine self-administration would increase calreticulin mRNA and protein. Male mice were trained to self-administer cocaine for 14 d and vHPC tissue punches were taken 1 d after the last administration (as in Fig. 1a). Cocaine self-administration significantly upregulated *Calr* mRNA in vHPC with a 2.73 fold change increase in cocaine compared to saline self-administering mice (Fig. 5d). From a separate cohort of cocaine and saline self-administering male mice, vHPC tissue was processed for total calreticulin protein (Fig. 5e, Suppl. Fig. 5). Cocaine significantly increased calreticulin protein in vHPC. We next sought to identify whether cocaine enhances ΔFosB binding to the *Calr* gene. We used the same experimenter-administration regimen (as Fig. 1f) in naïve male mice, and tissue was processed for ChIP-PCR. We found that cocaine treatment robustly enhanced ΔFosB binding to the *Calr* promoter region (Fig. 5f). Interestingly, we did not see significant ΔFosB binding to the *Calr* promoter in saline treated mice when compared to IgG controls, suggesting that this mechanism is driven by cocaine induction of ΔFosB.

We next sought to determine whether calreticulin is induced by cocaine in vHPC-NAc neurons. To address this, we used our circuit-tagging approach and treated mice with experimenter-administered cocaine (20 mg/kg; IP) for 10 d (Fig. 1f). vHPC tissue was taken 1 d after the last injection for immunohistochemistry staining of calreticulin. Calreticulin is highly conserved and has abundant expression across tissue and cell types (Krause & Michalak, 1997; M. Michalak, Milner, Burns, & Opas, 1992). Therefore, we quantified calreticulin staining intensity for each GFP+ vHPC-NAc neuron as a semi-quantitative measure of its expression in this circuit. We found that cocaine increases calreticulin intensity in vHPC-NAc neurons (Fig. 5g-h). Specifically, we observed a significant increase in calreticulin intensity across total cells. Collectively, these findings validate our TRAP-seq results and demonstrate that cocaine induction of ΔFosB drives calreticulin expression in vHPC-NAc neurons.

### Calreticulin mediates cocaine changes in vHPC-NAc excitability and behavior

The role of calreticulin in neurons is poorly understood, although there is evidence suggesting it plays a role in neuronal development (Hsu et al., 2005; Kotian et al., 2019; Qiu & Michalak, 2009; Shih et al., 2012), recovery from nerve injury (Kotian et al., 2019; Y. M. Lee, Park, Chung, & Oh, 2003; Pacheco, Merianda, Twiss, & Gallo, 2020), and memory (Kennedy, Kuhl, Barzilai, Sweatt, & Kandel, 1992; Kotian et al., 2019; Stoltzfus, Horton, & Grotewiel, 2003). Calreticulin’s role in neuronal excitability has not been investigated, although there is interesting evidence that calreticulin and neuronal excitability may be loosely linked to its role in ER Ca2+ homeostasis and chaperone functions (Saxena et al., 2013; Stutzmann & Mattson, 2011). Furthermore, calreticulin has been linked to cardiac excitability and development (Méry et al., 2005; Kimitoshi Nakamura et al., 2001). We therefore sought to determine whether calreticulin may also play a role in neuronal excitability. Our first step was to determine whether calreticulin upregulation was sufficient to decrease vHPC excitability. To address this question, we used viral overexpression of calreticulin in vHPC (AAV2-CMV-mCalr-GFP; or AAV2-CMV-GFP as control) in male C57Bl/6J mice. Approximately 4-6 weeks later, we assessed excitability using *ex vivo* slice electrophysiology in GFP-labeled calreticulin-overexpressing vCA1 neurons (Fig. 6a). Calreticulin overexpression reduced the excitability of vHPC neurons compared to control vHPC neurons (Fig. 6c-d). Specifically, we observed a calreticulin overexpression-induced decrease in spike frequency (Fig. 6b) and the total number of spikes (Fig. 6c), and a calreticulin overexpression-induced increase in rheobase (Fig. 6d). This suggests that calreticulin is sufficient to decrease vHPC excitability in the absence of cocaine or ΔFosB.

**Figure 6.**
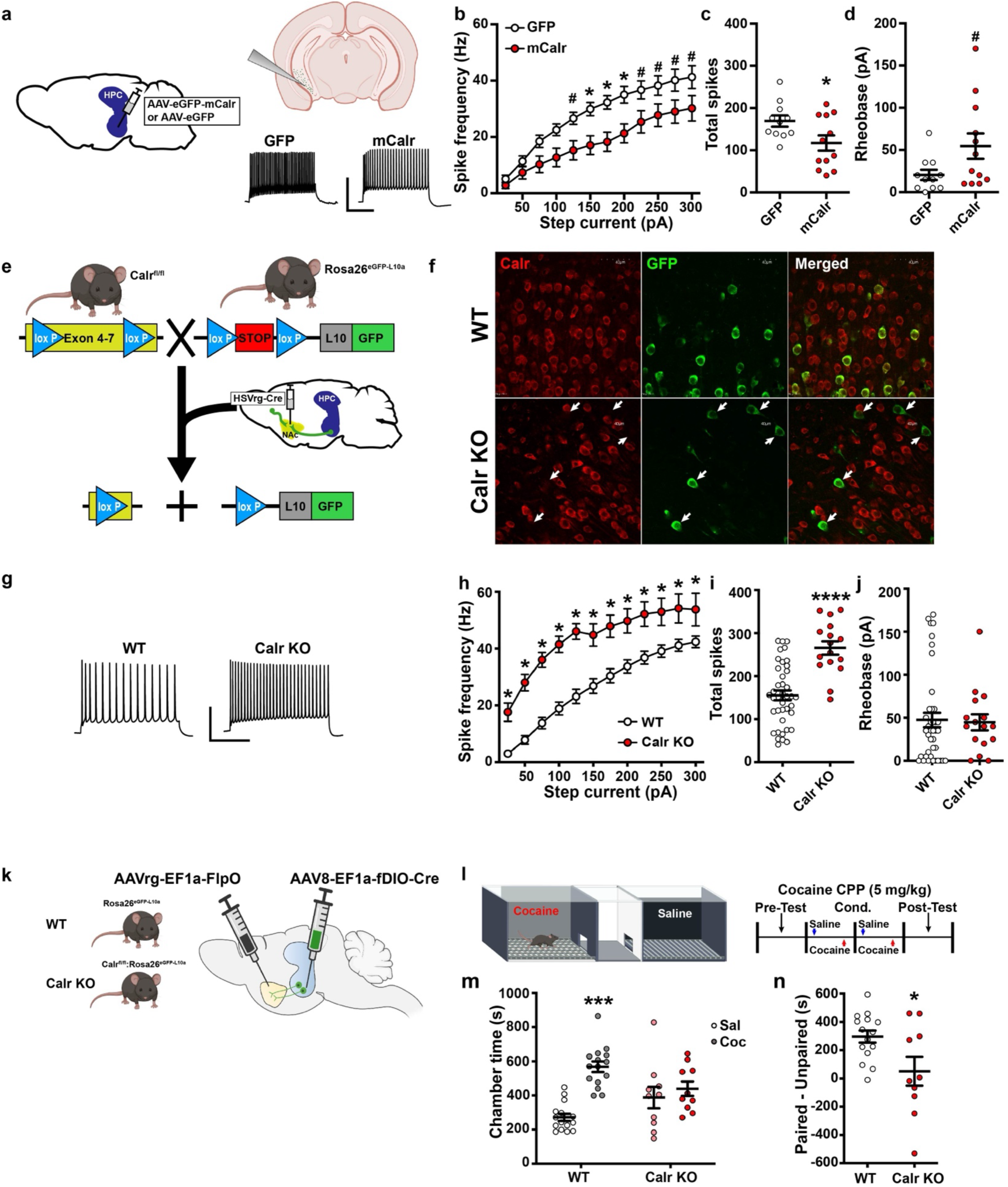
Calreticulin mediates ventral hippocampal excitability and behavior. **a)** Illustration of approach for overexpression of calreticulin on vHPC excitability, *ex vivo* patch clamp, and representative spike traces (300 pA; scale bar 50 pA, 200 ms). Male mice received AAV vector overexpressing calreticulin (mCalr, AAV2-CMV-mCalr-eGFP; or GFP, AAV2-CMV-eGFP as a control) into vHPC. 3-4 weeks later vCA1 neurons (n=11 cells/group; n=3 mice/group) were patched onto to assess excitability. **b)** mCalr overexpression reduced vCA1 spike frequency across depolarizing steps of current [Two-way RM ANOVA: ME of Group F (1, 20) = 7.600, P=0.0122; CurrentXGroup F (11, 220) = 1.935, P=0.0363], with significant decreases at current steps of 150-200 pA (Holm-Sidak *P<0.05), and a trend towards significant decreases at 225-300 pA (Holm-Sidak ^#^P<0.10). **c)** mCalr overexpression reduced the total number of spikes [Independent samples t-test: t(21)=2.298, *P=0.0319]. **d)** There was a trend towards a significant increase in rheobase by mCalr overexpression [Independent samples t-test: t(21)=2.041, ^#^P=0.0540]. **e)** Modified circuit-tagging approach to drive GFP and KO calreticulin in vHPC-NAc neurons. Male mice with crossed Floxed Calr and L10-GFP (KO; *Calr^fl/fl^*:*Rosa26^eGFP-L10a^*) or L10-GFP only (WT; *Rosa26^eGFP-L10a^*) received retrograde HSV vector expressing Cre (HSVrg-hEf1ɑ-Cre) into nucleus accumbens to drive GFP and Calr KO in projections to accumbens, including vHPC-NAc projections. **f)** Representative 20x coronal images in WT (top) and Calr KO (bottom) mice. Calreticulin stained neurons (red, left), GFP (green, middle). White arrows denote GFP+ neurons with a noted lack of calreticulin expression in Calr KO mice. **g)** Representative spike traces at 200 pA in vCA1-NAc neurons in WT (n=6 mice; n=40 cells) and Calr KO (n=4 mice; n=16 cells). Scale bar 50 pA, 200 ms. **h)** Calr KO significantly reduces vCA1-NAc neuron spike frequency across depolarizing steps of current [Two-way RM ANOVA: ME of Group F (1, 54) = 28.99, P<0.0001; CurrentXGroup F (11, 594) = 2.100, P=0.0187; Holm-Sidak *P<0.05]. **i)** Calr KO significantly increased total number of spikes compared to WT [Independent samples t-test: t(54)=5.409, ****P<0.0001]. **j)** There were no differences between groups on rheobase. **k)** Intersectional virus strategy to selectively knockout *Calr* in vCA1-NAc neurons. **l)** Cocaine CPP procedure. Male controls (WT; n=15) or Calr KO (Calr KO; n=10) mice underwent cocaine CPP in a 3-chamber box for twice-daily conditioning sessions of saline and cocaine (5 mg/kg; IP) followed by a drug-free post-test. **m)** Saline-paired (Sal) and cocaine-paired (Coc) chamber time during the post-test for WT and Calr KO mice. WT mice spent significantly more time in the Coc chamber compared to the Sal chamber, however Calr KO had no differences in time spent between chambers [Two-way RM ANOVA: ChamberXGroup: F (1, 23) = 6.311, P=0.0195; Holm-Sidak ***P=0.0002]. **d)** Cocaine-paired and cocaine-unpaired (i.e., saline) chamber difference in WT and Calr KO mice. Calr had significantly reduced paired-unpaired time compared to WT [Independent samples t-test: t(23)=2.512, *P=0.0195].

We next sought to determine whether calreticulin knockout in vHPC-NAc would alter the excitability of vCA1-NAc neurons. We received a generous donation of floxed calreticulin mice (*Calr^fl/f^*^l^) and crossed these mice with our *Rosa26^eGFP-L10a^* strain to produce a crossed line (*Calr^fl/f^*^l^: *Rosa26^eGFP-L10a^*). We used our circuit-tagging approach (Fig. 6e) to both drive GFP expression and knockout calreticulin (Calr KO) in all of the NAc-projecting neurons, including vHPC-NAc neurons. We performed immunohistochemistry for GFP and calreticulin in vHPC tissue slices from WT (*Rosa26^eGFP-L10a^*) and Calr KO male mice. We confirmed that Calr KO in vHPC-NAc neurons decreased calreticulin staining intensity (Fig. 6f-g). In a separate cohort of WT and Calr KO male mice we assessed the effects of Calr KO on vCA1-NAc excitability. We used our circuit-tagging approach (Fig. 6e) and measured excitability in vCA1-NAc neurons of WT and Calr KO mice. We observed that knockout of calreticulin increased the excitability of vCA1-NAc neurons (Fig. 6f-g), with no change in rheobase (Fig. 6h). These findings suggest that cocaine induction of calreticulin is a primary driver for cocaine-dependent changes in vHPC-NAc excitability.

Calreticulin changes in vHPC-NAc excitability may be a potential mediator of cocaine behavior, however few studies have identified the role of calreticulin in neurons and its role in behavior. We sought to determine the role of calreticulin in cocaine reward based on our observation that FosB knockout impairs cocaine reward (Fig. 2). We employed an intersectional viral strategy in crossed *Calr^fl/fl^: Rosa26^eGFP-L10a^*mice to KO calreticulin in only vHPC-NAc neurons (Fig. 6k). Briefly, male WT (*Rosa26^eGFP-L10a^*) and floxed calreticulin mice (*Calr^fl/fl^: Rosa26^eGFP-L10a^*) received viral infusions of AAVrg-Ef1a-FlpO into NAc and AAV8-EF1a-fDIO-Cre into vHPC. WT and Calr KO were then tested for cocaine conditioned place preference (Fig. 6l). Circuit-specific Calr KO in vHPC-NAc neurons reduced the preference for a context paired with cocaine (Fig. 6m-n). WT mice spent more time in a cocaine-paired chamber compared to a saline-paired chamber, however Calr KO mice spent a similar amount of time in both chambers. These data suggest that vHPC-NAc calreticulin is necessary for the expression of cocaine reward and support our overall model that cocaine induces ΔFosB to increase expression of calreticulin, reducing membrane excitability of vHPC-NAc neurons and driving cocaine reward and seeking.

## Discussion

It has been well established that cocaine produces plasticity at vHPC synapses onto NAc neurons (Boxer et al., 2023; Britt et al., 2012; Pascoli et al., 2014), yet these changes are largely postsynaptic in NAc MSNs. While there is no specific evidence that cocaine remodels presynaptic vHPC-NAc plasticity, this has been characterized in PFC synapses onto NAc MSNs (Suska, Lee, Huang, Dong, & Schlüter, 2013). Compelling evidence also suggests that cocaine conditioning and self-administration reshape the function of vHPC neurons, producing changes in excitatory transmission and dendritic spine morphology (Barr, Forster, & Unterwald, 2014; Barrientos et al., 2018; Caban Rivera et al., 2023; Keralapurath, Briggs, & Wagner, 2017; Keralapurath, Clark, Hammond, & Wagner, 2014; Preston & Wagner, 2022; Thompson, Swant, Gosnell, & Wagner, 2004). Here, we now demonstrate that cocaine remodels the intrinsic membrane excitability of vHPC-NAc neurons through induction of ΔFosB. The ΔFosB-mediated decrease in vHPC-NAc neuronal excitability mirrors prior findings that show that other psychostimulants reduce excitability in vSub (Cooper et al., 2003) and our own data that ΔFosB regulates dorsal CA1 (Eagle et al., 2018) and vCA1 excitability (Eagle et al., 2020). The mechanisms by which ΔFosB reduces excitability were previously unclear, however our analyses of gene expression in vHPC-NAc suggested both direct effects, e.g. ion channel expression, and indirect effects via signaling or downstream transcription factors. This lead to our in-depth investigation of the ER chaperone protein, calreticulin. In the future, it will be important to determine the time course by which ΔFosB reshapes the function of hippocampal neuron excitability, and how this impacts cellular and network function.

ΔFosB has a well-established role in nucleus accumbens and is regulated there by stress, natural rewards, and drug exposure (Nestler, 2015; Nestler, Barrot, & Self, 2001; Robison & Nestler, 2022), but its role in HPC is less clear. Previous studies showed that ΔFosB expression is induced in HPC (J. Chen et al., 1995; Eagle et al., 2015; Gajewski et al., 2019; Perrotti et al., 2008), and we demonstrated that cocaine induces ΔFosB in vHPC neurons (Gajewski et al., 2019). ΔFosB is a chronic activity-dependent transcription factor, therefore its induction regulates gene expression that likely leads to indelible changes in plasticity. Indeed, we observed a cocaine-induced change in vHPC-NAc gene expression that was, in part, ΔFosB-dependent.

We also observed that vHPC-NAc ΔFosB is necessary for cocaine reward and seeking. This is in line with our prior findings that cocaine exposure epigenetically modifies the *FosB* gene to enable ΔFosB expression in vHPC and drive cocaine reward (Gajewski et al., 2019) and that ΔFosB in dHPC is necessary for hippocampal-dependent learning and memory (Eagle et al., 2015). ΔFosB in nucleus accumbens is similarly regulated by cocaine and drives cocaine responses (Colby, Whisler, Steffen, Nestler, & Self, 2003; Kelz et al., 1999; McClung & Nestler, 2003; Robison et al., 2013; Yeh et al., 2023). However, its role in hippocampal-dependent regulation of cocaine behavior is not as well understood. It has been hypothesized that ventral hippocampus is necessary for linking external stimuli, e.g. cues, contexts, etc., with internal drives, such as the drive to seek and consume cocaine (V. S. Turner et al., 2022). This may require associative memory formation of cues or contexts associated with cocaine and other drugs and the underlying plasticity required for these phenomena. Indeed, ventral hippocampus projections to nucleus accumbens are necessary and sufficient for drug seeking (Bossert et al., 2016; Bossert & Stern, 2012; Britt et al., 2012; Caban Rivera et al., 2023; Lasseter et al., 2010; Rogers & See, 2007; Vorel et al., 2001), and the present work suggests that ΔFosB may therefore act as a molecular switch for gene expression in this circuit underlying a cocaine memory.

It is unclear how a ΔFosB expression-dependent decrease in vHPC-NAc neuron excitability affects NAc MSNs. The activity of glutamatergic afferent synapses onto NAc MSNs regulates synaptic plasticity (Kalivas, 2009; B. D. Turner, Kashima, Manz, Grueter, & Grueter, 2018; van Huijstee & Mansvelder, 2015), which is particularly affected by drugs of abuse including cocaine, ultimately leading to increased relapse (Lüscher & Malenka, 2011; Pascoli et al., 2014; Russo et al., 2010; Wolf, 2010; Wolf & Ferrario, 2010). However, cocaine can also produce homeostatic plasticity changes in NAc MSNs (Delint-Ramirez et al., 2020; Dong et al., 2006; Ishikawa et al., 2009; B. R. Lee & Dong, 2011; X. Sun & Wolf, 2009; J. Wang et al., 2018). It may be that a decrease in vHPC-NAc excitability leads to changes in feedforward inhibition in NAc MSNs (Scudder, Baimel, Macdonald, & Carter, 2018; Yu et al., 2017). These changes could also be dependent on whether vHPC synapses onto D1 or D2 MSNs. Relapse to cocaine seeking produces increased synaptic plasticity at vHPC synapses onto D1 MSNs (Lüscher & Malenka, 2011), and cocaine produces feedforward inhibition largely at D1 MSNs (Scudder et al., 2018), however recent evidence shows that cocaine biases plasticity toward vSub neuron synapses onto D2 MSNs (Boxer et al., 2023). We have also observed that ΔFosB expression in vHPC-NAc neurons mediates stress susceptibility and vHPC-NAc excitability (Eagle et al., 2020). Stress and drugs may produce alterations in vHPC that lead to synergistic or opposing downstream changes in the NAc, an idea supported by previous work showing that both cocaine and stress remodel vHPC synapses onto NAc (LeGates et al., 2018). These feedforward changes may have widespread impacts on mood and reward function, such as stress susceptibility and susceptibility to drug seeking.

The mechanism for calreticulin changes in vHPC excitability and cocaine behavior remains to be explored. Calreticulin is a high-capacity, low-affinity, Ca2+-binding chaperone protein that, along with calnexin, regulates endoplasmic reticulum (ER) Ca2+ storage and protein folding (Marek Michalak et al., 2009). Calreticulin’s low affinity for Ca2+ is important in maintaining large concentrations of free Ca2+ in the ER available for rapid release, with calreticulin accounting for ∼50% of the ER free Ca2+ capacity (Arnaudeau et al., 2002; Bastianutto et al., 1995; Krause & Michalak, 1997; Marek Michalak et al., 2009; K. Nakamura et al., 2001; W.-A. Wang, Groenendyk, & Michalak, 2012). While cocaine effects on ER Ca2+ release have not been investigated, cocaine reduces ER store-operated Ca2+ entry (Brailoiu et al., 2016). It is quite possible that calreticulin is a mechanism for cocaine’s effects on ER regulation of intrinsic neuronal excitability, and the ER is a vital organelle in neurons for maintaining calcium (Ca2+) homeostasis. The ER can enhance intracellular Ca2+-mediated signaling, synaptic plasticity, and vesicle release (Bardo, Cavazzini, & Emptage, 2006; Maggio & Vlachos, 2014; Mattson et al., 2000; Segal & Korkotian, 2014; Verkhratsky, 2005). ER Ca2+ stores are replenished primarily through SERCA (sarco-endoplasmic reticulum Ca2+ ATPase) and CRAC (Ca2+ release activated channel), whereas Ca2+ release is primarily conducted through ryanodine receptors (RyR) and inositol triphosphate receptors (IP3R). Release of Ca2+ from ER stores can be initiated by a variety of signaling events, including changes in cytosolic Ca2+ and metabotropic receptor signaling. However, the role of ER Ca2+ release in regulating neuronal physiology (i.e. excitability) is not well understood, though there is a compelling argument from the ER’s role in synaptic plasticity (Bardo et al., 2006; Maggio & Vlachos, 2014; Mattson et al., 2000; Segal & Korkotian, 2014) that the ER may also alter long-term excitability. Therefore, cocaine induction of calreticulin through ΔFosB may produce changes in ER that lead to decreased vHPC excitability and subsequent cocaine seeking.

One potential drawback to the generalizability of these findings is that we only included male mice in our studies. This was primarily due to our own observations that vHPC-NAc neuron excitability in females (in the absence of cocaine or any other manipulation) is quite different than males (Williams et al., 2020). This difference is driven by adult testosterone and reduces stress-driven anhedonia in male mice compared to females. There have been multiple findings showing that female mice are more likely to escalate drug self-administration, reinstate at greater rates, and are more motivated for drugs (Becker & Koob, 2016; Bobzean, DeNobrega, & Perrotti, 2014; Calipari et al., 2017; Doncheck et al., 2020; Festa et al., 2004; Fuchs, Evans, Mehta, Case, & See, 2005; Hu, Crombag, Robinson, & Becker, 2004; Lynch & Carroll, 1999; Nicolas et al., 2019; Perry, Westenbroek, Jagannathan, & Becker, 2015; Quigley et al., 2021; Russo et al., 2003; Song, Yang, Peckham, & Becker, 2019; Swalve, Smethells, & Carroll, 2016; Takashima et al., 2018; Yousuf et al., 2019). Sex differences also exist in hippocampal structure and function (Koss & Frick, 2017; Smith, Jones, & Wilson, 2002; van Eijk et al., 2020; Yagi & Galea, 2019), supporting clinical evidence for sex differences in addiction (Becker, 2016; Becker, McClellan, & Reed, 2016; Becker, McClellan, & Reed, 2017; Bobzean et al., 2014; Potenza et al., 2012). Thus, an additional potential mediator for cocaine changes in gene expression could be sex-dependent effects on vHPC-NAc excitability, and that will certainly be a focus of future studies in our group and others.

## Methods

### Animals

All experiments were approved by the Institutional Animal Care and Use Committee at Michigan State University in accordance with AAALAC. Male C57Bl/6J mice (3–5/cage, 7–8-week old upon arrival from Jackson Labs) were allowed at least 5 d to acclimate to the facility prior to any experimental procedures. The floxed FosB mouse strain (*FosB^fl/fl^*) was a generous gift from the laboratory of Dr. Eric Nestler at the Icahn School of Medicine at Mount Sinai. The *Rosa26^eGFP-L10a^* mice were a generous gift from the laboratory of Dr. Gina Leinninger at Michigan State University. The floxed calreticulin mouse strain (*Calr^fl/fl^*) was originally created by Dr. Masahito Ikawa and was a generous gift from the laboratory of Dr. Marek Michalak. Unless otherwise stated, all mice were group housed in a 12:12 h light/dark cycle with *ad libitum* food and water. Temperature (22 °C) and humidity (50–55%) were held constant in animal housing and behavioral testing rooms.

### Stereotaxic surgery and viral vectors

Stereotaxic surgery was conducted as previously described (Eagle et al., 2020). For circuit-tagging experiments (Fig. 1f), retrograde Cre vector (HSV-hEf1α-Cre; 0.5 μL; Gene Delivery Core, Massachusetts General Hospital) was infused into NAc (+1.6 AP, ±1.5ML, −4.4DV relative to bregma, 10° angle). For CRISPR experiments, retrograde Cas9 vector (HSV-hEf1α-LS1L-myc-Cas9; 0.5 μL; Gene Delivery Core, Massachusetts General Hospital) was infused into NAc. After 3 weeks, control vector (HSV-IE4/5-TB-eYFP-CMV-IRES-Cre; Gene Delivery Core, Massachusetts General Hospital) or FosB gRNA vector (HSV-IE4/5-TB-FosB gRNA-CMV-eYFP-IRES-CRE; Gene Delivery Core, Massachusetts General Hospital) was infused into the ventral CA1 region of vHPC (vCA1; −3.4AP, ±3.2ML, −4.8DV relative to bregma, 3° angle; 0.5 μL). For DREADD experiments, retrograde Cre vector was infused into NAc and control vector (AAV2-CMV-GFP; 0.5 μL; UNC Vector Core) or DREADD Gq vector [AAV2-hSyn-DIO-hM3D(Gq)-mCherry; 0.5 μL; AddGene] was infused into the vCA1. For calreticulin overexpression experiments, mCalr vector (AAV2-CMV-mCalr-SV40p-eGFP; 0.5 μL; Vector BioLabs) was infused into vCA1. For circuit-specific calreticulin knockout experiments, retrograde FlpO vector (AAVrg-Ef1a-FlpO; 0.5 μL; Addgene) was infused into NAc and a Flp-dependent Cre virus (AAV8-EF1a-fDIO-Cre0.5 μL; Addgene) was infused into vCA1. All behavioral and electrophysiological procedures commenced 3-6 weeks following surgeries.

### Behavioral testing

Behavioral data were collected using an IR-CCD camera (Panasonic) and analyzed using automated video tracking software (CleverSys). Animals were transported to the behavioral testing rooms in their home cages and allowed 30 min to habituate to the room.

#### Experimenter-administered cocaine

Cocaine hydrochloride (dissolved in 0.9% sterile saline) was injected IP into mice daily in their homecages for 10 d.

#### Cocaine self-administration

Cocaine self-administration was conducted as previously described (Doyle et al., 2021). Briefly, mice were trained and tested for seeking in a mouse operant chamber equipped for self-administration (Med-Associates, Inc.), including an automated syringe pump, commutator and infusion tubing, magazine, house light, and left and right nosepokes with cue lights within the nosepoke hole. Mice received jugular catheter surgeries, and 4 d following surgery were trained *de novo* to self-administer IV infusions of cocaine or saline (0.5 mg/kg/infusion) across 14 daily 2 h sessions. House and cue lights remained on until a successful FR1 ratio nosepoke response in the active nosepoke hole (randomly assigned left or right). This was immediately followed by an infusion (including the sound of the syringe pump) and a 20 s timeout period of the house and cue lights. Nosepoke responses on the inactive nosepoke had no consequences. Responses were limited to 75 responses (37.5 mg/kg total cocaine) in each session. For the abstinence-induced seeking test, animals were left in their homecages for 6 d after the last administration session and then were tested again in the same operant chambers. The seeking test was the same as administration session except: 1) nosepoke responses on the active nosepoke resulted in no infusion but the sound of the pump; and 2) session duration was 1 h.

### Cocaine conditioned place preference (CPP)

CPP was conducted as previously described (Gajewski et al., 2019). Briefly, mice were tested for cocaine CPP in a 3-chamber CPP box (San Diego Instruments). On day 1, mice received a pre-test (no cocaine or saline) during which they were allowed to explore the entire box for 15-20 min. On days 2-3, mice received injections of saline paired with one chamber in the morning for 30 min and injections of cocaine (5 mg/kg; IP) paired with the opposite chamber for 30 min in the afternoon (chamber counterbalanced by group). On day 4, mice were again tested in a post-test (no cocaine or saline) and allowed to freely explore the entire box.

#### Chemogenetic experiments

Gq (and control) mice were treated daily with clozapine-N-oxide (CNO; 0.3 mg/kg) dissolved in 5% DMSO. Mice received twice-daily injections of CNO (in the morning and afternoon). Prior to CPP, CNO was injected and mice were left in their home cages. During CPP, CNO was injected 20-30 min prior to testing and conditioning.

#### Immunohistochemistry

Immunofluorescent analysis was conducted as previously described (Eagle et al., 2020). Mice were transcardially perfused with cold PBS, followed by 10% formalin. Brains from all immunostaining experiments were postfixed 24 h in 10% formalin, cryopreserved in 30% sucrose, and sliced frozen on an SM2010R microtome (Leica) into 35 μm sections. Immunohistochemistry was performed using primary antibodies against FosB (ab11959; 1:1000; Abcam), calreticulin (1:2000; ab92516, Abcam), and GFP (ab5450; 1:1000-1:5000; Abcam); and secondary antibodies (1:200; Jackson Immunoresearch) conjugated to fluorescent markers (AlexaFluor 488; Cy3; Cy5). Fluorescent images were visualized on an Olympus FluoView 1000 filter-based laser scanning confocal microscope or a Nikon Eclipse Ni-U Upright Fluorescent Microscope. Cell counts and intensity of signal in individual cells was quantified using ImageJ (NIH) software by an experimenter blinded to conditions.

#### RT-PCR

RNA extraction and RT-PCR was conducted as previously described (Eagle et al., 2020). Mice were sacrificed and fresh brains were immediately dissected into 1 mm coronal sections. vHPC tissue was collected using 14-gauge biopsy punches guided by a fluorescent dissecting microscope (Leica) and stored at −80 °C until processing. RNA was isolated from vHPC tissue using TriZol (Invitrogen) homogenization and chloroform layer separation. The clear RNA layer was then processed (RNAeasy MicroKit, Qiagen #74004) and analyzed with NanoDrop. A volume of 10 μL of RNA was reverse transcribed to cDNA (High Capactiy cDNA Reverse Transcription Kits Applied BioSystens #4368814). Prior to qPCR, cDNA was diluted to 200 μL. The reaction mixture consisted of 10 μL PowerSYBR Green PCR Master Mix (Applied Biosystems; #436759), 2 μL each of forward and reverse primers and water, and 4 μL cDNA template. Samples were then heated to 95 °C for 10 min (Step 1) followed by 40 cycles of 95 °C for 15 s, 60 °C for 15 s, and 72 °C for 15 s (Step 2), and 95 °C for 15 s, 60 °C for 15 s, 65 °C for 5 s, and 95 °C for 5 s (Step 3). Analyses were carried out using the ΔΔC(t) method (Schmittgen et al., 2000). Samples were normalized to *Gapdh*.

### Calr

Forward: 5’ GAA TAC AAG GGC GAG TGG AA 3’ Reverse: 5’ GGG GGA GTA TTC AGG GTT GT 3’

#### Western blotting

Western blotting analysis was conducted as previously described (Gajewski et al., 2019). Mice were sacrificed and fresh brains were immediately dissected into 1 mm coronal sections. vHPC tissue was collected using 14-gauge biopsy punches guided by a fluorescent dissecting microscope (Leica) and stored at −80 °C until processing. vHPC tissue punch samples were processed for SDS-PAGE and transferred to PVDF membranes for Western blotting with chemiluminescence. Blots were probed with calreticulin (1:1000; ab92516, Abcam), and then stripped and reprobed for Gapdh (14C10; 1:20,000; 2118, Cell Signaling Technology). Protein was quantified relative to Gapdh using ImageJ (NIH) software.

#### Translating ribosomal affinity purification (TRAP) and cDNA library preparation

TRAP was conducted as previously described (Eagle et al., 2020). Three weeks following injection of retrograde HSV-Cre into NAc, Cre-dependent L10-GFP-expressing mice (*Rosa26^eGFP/L10a^*) were sacrificed and fresh brains were immediately dissected into 1 mm coronal sections. vHPC tissue was collected using 14-gauge biopsy punches guided by a fluorescent dissecting microscope (Leica) and stored at −80 °C until processing (n = 19-20 punches/group, 3–4 mice pooled per group). Polyribosome-associated RNA was affinity purified by homogenizing vHPC tissue in ice-cold tissue-lysis buffer (20mM HEPES [pH 7.4], 150mM KCl, 10mM MgCl2, 0.5mM dithiothreitol, 100 μg/ml cycloheximide, protease inhibitors, and recombinant RNase inhibitors) using a motor-driven Teflon glass homogenizer. Homogenates were centrifuged for 10 min at 2000×g (4 °C), supernatant was supplemented with 1% NP-40 (AG Scientific, #P1505) and 30mM DHPC (Avanti Polar Lipids, #850306P), and centrifuged again for 10 min at 20,000 × g (4 °C). Supernatant was collected and incubated with Streptavidin MyOne T1 Dynabeads (Invitrogen, #65601) that were coated with anti-GFP antibodies (Memorial Sloan-Kettering Monoclonal Antibody Facility; clone names: Htz-GFP-19F7 and Htz-GFP-19C8, 50 μg per antibody per sample) using recombinant biotinylated Protein L (Thermo Fisher Scientific, #29997) for 16–18 h on a rotator (4 °C) in low salt buffer (20mM HEPES [pH 7.4], 350mM KCl, 1% NP-40, 0.5mM dithiothreitol, 100 μg/ml cycloheximide). Beads were isolated and washed with high-salt buffer (20mM HEPES [pH 7.4], 350mM KCl, 1% NP-40, 0.5mM dithiothreitol, 100 μg/ml cycloheximide) and RNA was purified using the RNeasy MicroKit (Qiagen, #74004). In order to increase yield, each RNA sample was initially passed through the Qiagen MinElute™ column three times. Following purification, RNA was quantified using a Qubit fluorometer (Invitrogen) and RNA quality was analyzed using a 4200 Agilent Tapestation (Agilent Technologies). cDNA libraries from 5 ng total RNA were prepared using the SMARTer® Stranded Total RNA-Seq Kit (Takara Bio USA, #635005), according to manufacturer’s instructions. cDNA libraries were pooled following Qubit measurement and TapeStation analysis, with a final concentration of 4 nM.

#### Sequencing

Sequencing was performed at the Icahn School of Medicine at Mount Sinai Genomics Core Facility (https://icahn.mssm.edu/research/genomics/core-facility). Raw sequencing reads from mice were mapped to mm10 using HISAT2. Counts of reads mapping to genes were obtained using featureCounts software of Subread package against Ensembl v90 annotation. Differential expression was conducted using the DESeq2 package (Love, Huber, & Anders, 2014). Sequencing data has been deposited into the GEO database (Accession number: GSE281142).

### Chromatin Immunoprecipitation (ChIP-PCR)

Brain tissues were suspended in phosphate-buffered saline and disrupted by passing them through 25-gauge needles. The disrupted tissues were then fixed with 2 mM ethylene glycol bis(succinimidylsuccinate) (Thermo Scientific) for 1 hour, followed by treatment with 1% formaldehyde for 15 minutes and 0.125 M glycine for 5 minutes to quench the reaction. The cells were lysed in a solution containing 1% SDS, 10 mM EDTA, and 50 mM Tris–HCl (pH 8.0). The DNA was fragmented into approximately 200–400 base pair lengths using sonication (Branson Sonifier 450). Immunoprecipitation was carried out overnight at 4 °C using 5 μl of rabbit polyclonal FosB antibody (2251S, lot 3, Cell Signaling Technology) or 2 μg of rabbit polyclonal IgG isotype (3900S, Cell Signaling Technology). Antibody-bound chromatin fragments were isolated with protein G plus/protein A agarose beads (MilliporeSigma), followed by washing and elution. After the immunoprecipitated and input chromatin fragments were reverse cross-linked, the DNA was extracted using phenol/chloroform and subsequently precipitated.

FosB occupancy at the *Calr* promoter was analyzed using quantitative real-time PCR on the C384 Real-Time System (Bio-Rad). The primer sequences used were GACCTACAGCTGTCCCTTTC (forward) and CCCATAGTGCGACCAATAGAA (reverse). Binding enrichment was quantified as the percentage of antibody (IgG isotype or FosB) immunoprecipitated DNA fragments relative to the input.

#### Electrophysiology

Whole-cell, *ex vivo* slice electrophysiology was conducted as previously described (Eagle et al., 2020; Eagle et al., 2018). All solutions were bubbled with 95% O2−5% CO2 throughout the procedure. Mice were anesthetized with isoflurane anesthesia and transcardially perfused with sucrose artificial cerebrospinal fluid (sucrose aCSF; in mM: 234 sucrose, 2.5 KCl, 1.25 NaH2PO4, 10 MgSO4, 0.5 CaCl, 26 NaHCO3, 11 glucose). Brains were rapidly removed, blocked, and placed in cold sucrose aCSF. Coronal sections (250 μM) containing vHPC were cut on a vibratome (Leica) and transferred to an incubation chamber containing aCSF (in mM: 126 NaCl, 2.5 KCl, 1.25 NaH2PO4, 2 MgCl, 2 CaCl, 26 NaHCO3, 10 glucose) held at 34 °C for 30 min before moving to aCSF at room temperature until used for recordings. Recordings were made from a submersion chamber perfused with aCSF (2 mL/min) held at 32 °C. Borosilicate glass electrodes (3–6MΩ) were filled with K-gluconate internal solution (in mM: 115 potassium gluconate, 20 KCl, 1.5 MgCl, 10 phosphocreatine-Tris, 2 MgATP, 0.5 Na3GTP; pH 7.2–7.4; 280–285 mOsm). GFP-positive cells in the ventral CA1 region of HPC were visualized using an Olympus BX51WI microscope using DIC infrared and epifluorescent illumination. Whole-cell patch-clamp recordings were made from transfected cells using a Multiclamp 700B amplifier and Digidata 1440A digitizer (Molecular Devices) and whole-cell junction potential was not corrected. Traces were sampled (10 kHz), filtered (10 kHz), and digitally stored. Cells with membrane potential more positive than −50 mV or series resistance >20MΩ were omitted from analysis. Rheobase was measured by giving brief (250 ms) depolarizing (0-300 pA, Δ5 pA steps) steps with 10 s step intervals. Elicited spike number was measured by issuing increasing depolarizing steps (0–300 pA, Δ25 pA steps, 500 ms) with 30 s step intervals. All electrophysiology recordings were made at approximately 30-32 °C by warming the aCSF line with a single inline heater (Warner Instruments).

#### Statistics and Reproducibility

We used one-way and two-way ANOVAs (including repeated measures or mixed effects as appropriate) followed by Holm-Sidak corrected *post hoc* comparisons in the case of significant omnibus effects, and independent samples t-tests. Alpha criterion was set to 0.05. To ensure reproducibility, separate cohorts were included in behavioral testing.

