## Supplemental Figures for "Cocaine, via ΔFosB, remodels gene expression and excitability in ventral hippocampus"

**Supplemental Figure 1. Cocaine or saline self-administration responses and accuracy.** Cocaine (0.5 mg/kg/infusion; IV; n=8) or saline (n=5) self-administration performance across 14 daily sessions in male mice. **a)** Nosepoke responses per min across cocaine or saline self-administration. Cocaine self-administering mice responded on the active nosepoke more than saline self-administering mice across sessions [Two-way RM ANOVA: ME of Group: $F(1, 11) = 19.54$, $P=0.0010$; ChamberXGroup: $F(13, 143) = 2.169$, $P=0.0137$], with significantly increased responding on days 6 & 11-14 (Holm-Sidak $^*P<0.05$). No differences were observed between groups in inactive nosepoke responses. **b)** Performance accuracy, i.e. % responding on the active lever compared to inactive lever, across cocaine or saline self-administration. Accuracy was overall decreased in saline self-administering mice compared to saline self-administering mice [Two-way RM ANOVA: ME of Group: $F(1, 11) = 9.229$, $P=0.0013$].

**Supplemental Figure 2. FosB knockout in vHPC-NAc effects on cocaine self-administration and forced abstinence-induced seeking responses and accuracy.** Cocaine (0.5 mg/kg/infusion; IV) self-administration and forced abstinence-induced seeking performance in control (Con; n=6) and CRISPR FosB KO (crKO; n=12) male mice. **a)** Nosepoke responses per min across 14 d of cocaine self-administration in Con and crKO mice. No differences were observed between groups in active or inactive nosepoke responses during self-administration. **b)** Nosepoke responses per min on a forced abstinence-induced seeking test at Day 21 (7 d after self-administration) in Con and crKO mice. crKO significantly reduced responding on the active nosepoke compared to Con mice [Independent samples t-test: $t(15)=2.171$, $^*P=0.0464$]. There were no differences between groups in inactive nosepoke responding (data not shown). **c)** Performance accuracy, i.e. % responding on the active lever compared to inactive lever, across cocaine self-administration in Con and crKO mice. There were no differences between groups in accuracy across self-administration. **d)** Accuracy during a forced abstinence-induced seeking test at Day 21 in Con and crKO mice. No differences in accuracy were observed between groups.

**Supplemental Figure 3. Cocaine self-administration behavior for electrophysiology studies.** Cocaine (Coc; 0.5 mg/kg/infusion; IV; n=7) or saline (Sal; n=6) self-administration performance across 14 daily sessions in male mice. **a)** Nosepoke responses per min across 14 d of cocaine self-administration in Sal and Coc mice. Cocaine self-administering mice responded on the active nosepoke more than saline self-administering mice across sessions [Two-way RM ANOVA: ME of Group: $F (1, 11) = 42.20$, $P<0.0001$; ChamberXGroup: $F (13, 143) = 2.672$, $P=0.0022$], with significantly increased responding on days 4, 9-12, & 14 (Holm-Sidak *$P<0.05$). No differences were observed between groups in inactive nosepoke responses. **b)** Infusions per min across 14 d of cocaine self-administration in Sal and Coc mice. Coc mice received more infusions compared to Sal mice [Two-way RM ANOVA: ME of Group: $F (1, 11) = 74.09$, $P<0.0001$; ChamberXGroup: $F (13, 143) = 4.368$, $P<0.0001$], with significantly more infusions on days 4-5 & 8-14 (Holm-Sidak *$P<0.05$). **c)** Performance accuracy, i.e. % responding on the active lever compared to inactive lever, across cocaine self-administration in Sal and Coc mice. Coc mice had significantly higher accuracy compared to Sal mice [Two-way RM ANOVA: ME of Group: $F (1, 11) = 16.53$, $P<0.0019$; ChamberXGroup: $F (13, 143) = 1.969$, $P=0.0274$], with significantly greater accuracy on days 5 & 13 (Holm-Sidak *$P<0.05$).

**Supplemental Figure 4. Subacute activation of vHPC-NAc does not impair cocaine CPP. a)** Viral strategy and experimental timeline. A retrograde HSV vector expressing Cre (HSVrg-hEf1α-Cre) was injected into nucleus accumbens and a Cre-dependent DREADD Gq viral vector (AAV2-hSyn-DIO-hM3Dq-mCherry) was injected into ventral hippocampus. Clozapine-N-oxide (CNO) or vehicle (Veh; 5% DMSO) was administered in separate groups 30 min before each testing and conditioning session **c)** Saline-paired (Sal) and cocaine-paired (Coc) chamber time during the post-test for Veh (n=8) and CNO (n=8) mice. Overall, both groups spent significantly more time in the Coc chamber compared to the Sal chamber [Two-way RM ANOVA: ME of Chamber: $F (1, 14) = 7.406$, $P=0.0165$]. **d)** Cocaine-paired and cocaine-unpaired chamber difference in Veh and CNO mice. There were no significant differences between groups.

**Supplemental Figure 5. Western blot images from vCA1 tissue of cocaine self-administering mice.** Western blot image of calreticulin in saline (S) and cocaine self-administering male mice (C) showing bands of around 55 kDa, close to the predicted molecular weight of 60 kDa.

### Supplemental Tables

**Supplemental Table 1. Cellular properties of vCA1 neurons in saline and cocaine self-administering male mice.**

| | MEASURES | | |
|---|---|---|---|
| **GROUPS:** | **RMP (mV)** | **Cm (pF)** | **Rm (MΩ)** |
| **Sal** | -67.8 ± 0.9 | 58.6 ±4.9 | 192.1 ±10.5 |
| **Coc** | -67.1 ±0.8 | 59.2 ±6.2 | 228.0 ±23.4 |

RMP = resting membrane potential; Cm = membrane capacitance; Rm = membrane resistance

**Supplemental Table 2. Cellular properties of vCA1-NAc neurons in saline- and cocaine-treated WT and FosB KO male mice.**

| | MEASURES | | |
|---|---|---|---|
| **GROUPS:** | **RMP (mV)** | **Cm (pF)** | **Rm (MΩ)** |
| **Sal** | -65.8 ± 1.1 | 79.2 ±6.0 | 223.0 ±29.4 |
| **Coc** | -66.2 ±1.2 | 72.0 ±4.7 | 249.2 ±25.3 |

RMP = resting membrane potential; Cm = membrane capacitance; Rm = membrane resistance

**Supplemental Table 3. Cellular properties of vCA1-NAc neurons in saline and cocaine self-administering WT male mice.**

| | MEASURES | | |
|---|---|---|---|
| **GROUPS:** | **RMP (mV)** | **Cm (pF)** | **Rm (MΩ)** |
| **WT-Sal** | -65.4 ± 1.0 | 43.8.2 ±2.0 | 130.7 ±16.7 |
| **WT-Coc** | -65.7 ±1.2 | 46.5 ±3.7 | 219.5 ±101.4 |
| **FosB KO-Coc** | -68.4 ±2.0 | *60.2 ±3.9** | *81.4 ±7.6** |

RMP = resting membrane potential; Cm = membrane capacitance; Rm = membrane resistance

**Supplemental Table 4. Cellular properties of vCA1 neurons in GFP and calreticulin overexpression (mCalr) male mice.**

| | MEASURES | | |
|---|---|---|---|
| **GROUPS:** | **RMP (mV)** | **Cm (pF)** | **Rm (MΩ)** |
| **GFP** | -65.0 ± 1.2 | 72.0 ±11.0 | 201.1 ±25.0 |
| **mCalr** | -67.1 ±1.2 | 73.9 ±6.9 | 204.1 ±26.3 |

RMP = resting membrane potential; Cm = membrane capacitance; Rm = membrane resistance

**Supplemental Table 5. Cellular properties of vCA1-NAc neurons in WT and Calreticulin KO male mice.**

| | MEASURES | | |
|---|---|---|---|
| **GROUPS:** | **RMP (mV)** | **Cm (pF)** | **Rm (MΩ)** |
| **WT** | -64.3 ± 1.1 | 106.6 ±5.4 | 192.7 ±24.8 |
| **Calr KO** | -65.9 ±1.5 | 95.3 ±11.8 | 163.8 ±18.6 |

RMP = resting membrane potential; Cm = membrane capacitance; Rm = membrane resistance

Supplemental Figure 1.

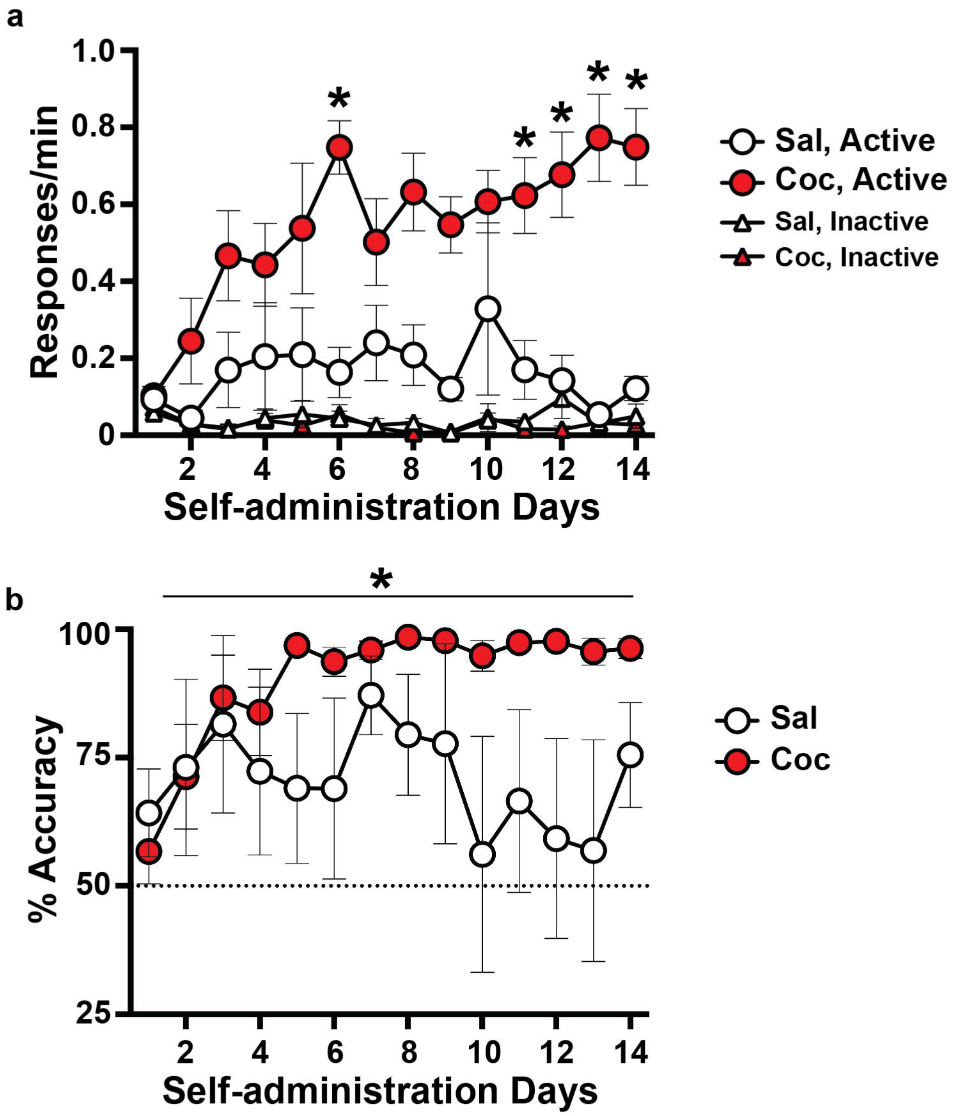

Supplemental Figure 2.

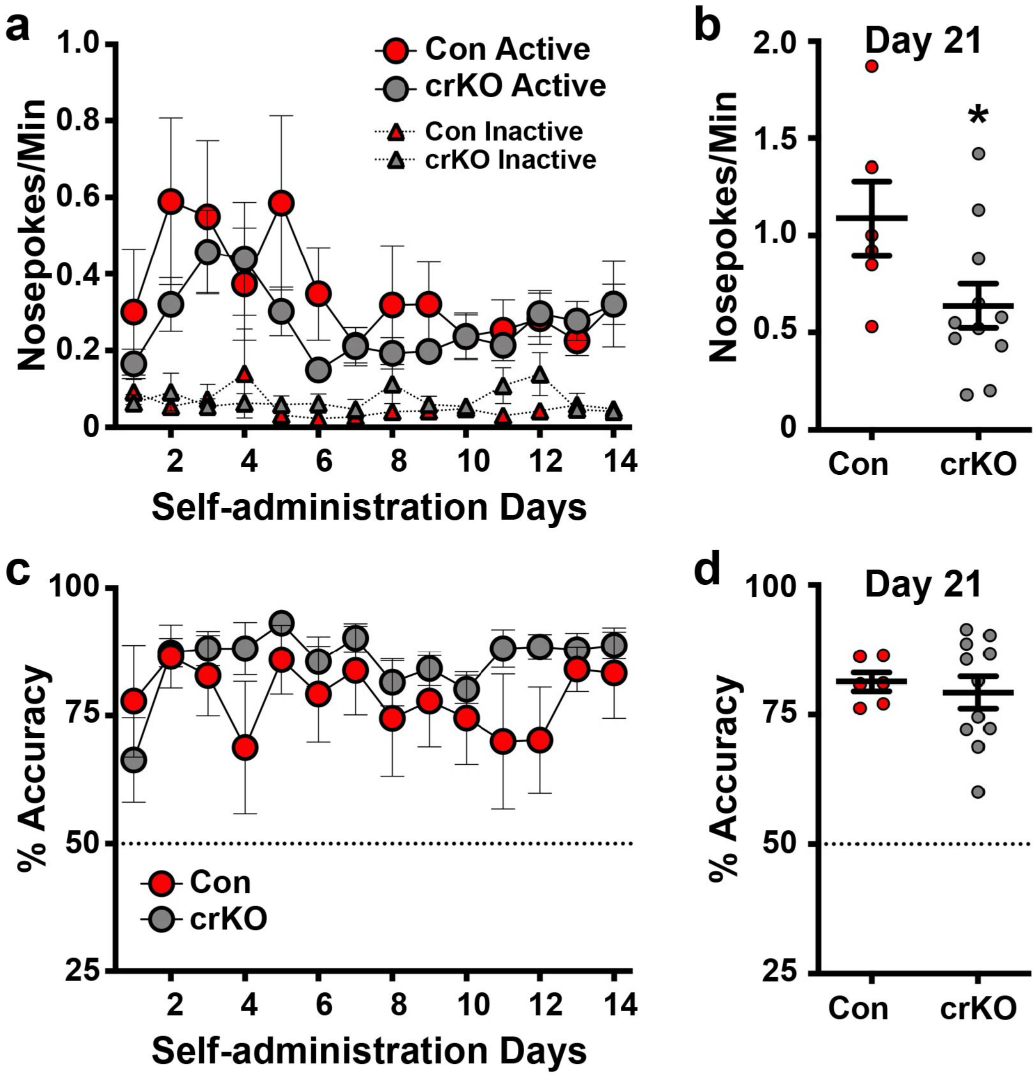

**Supplemental Figure 3.**

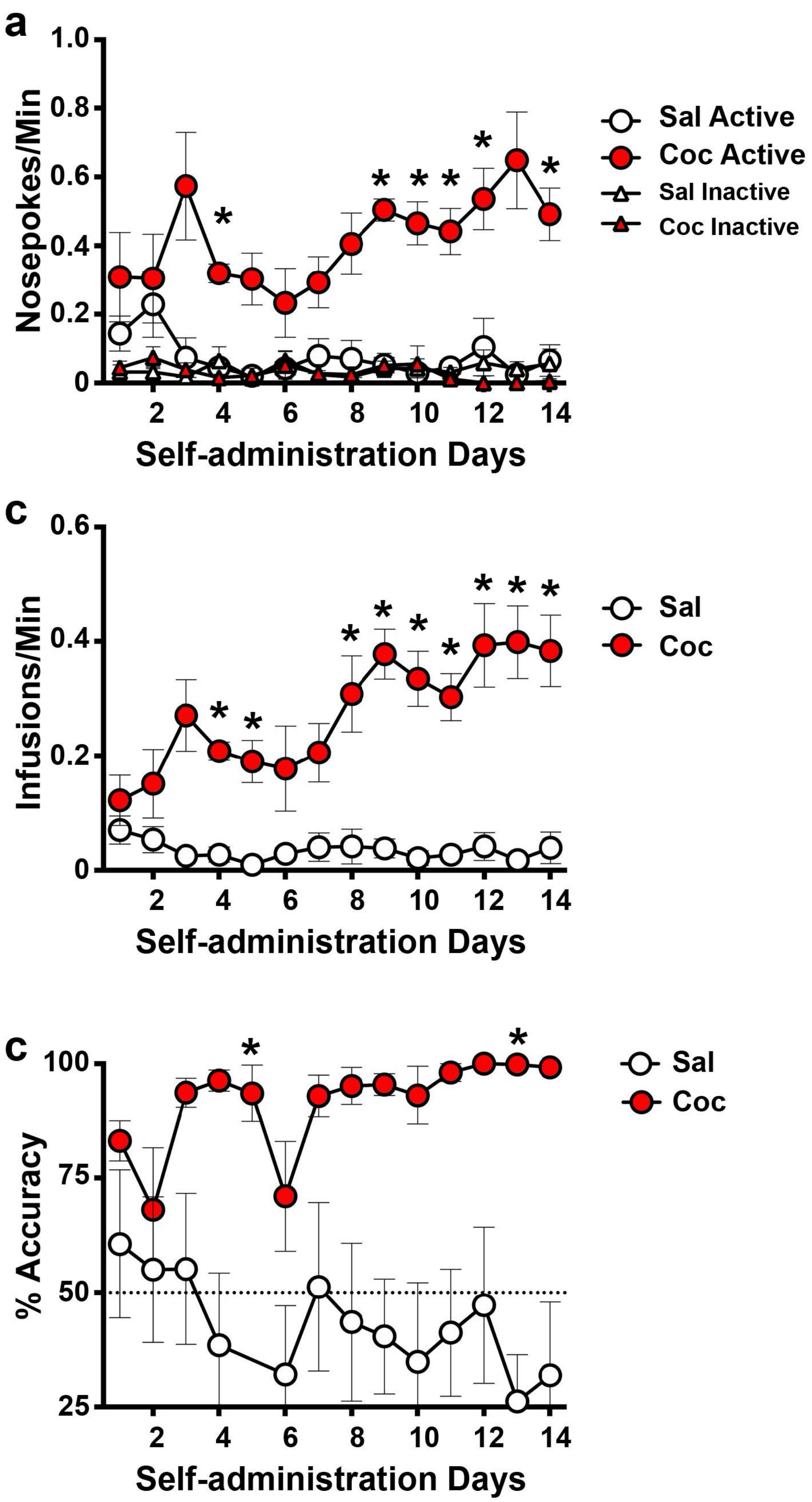

**Supplemental Figure 4.**

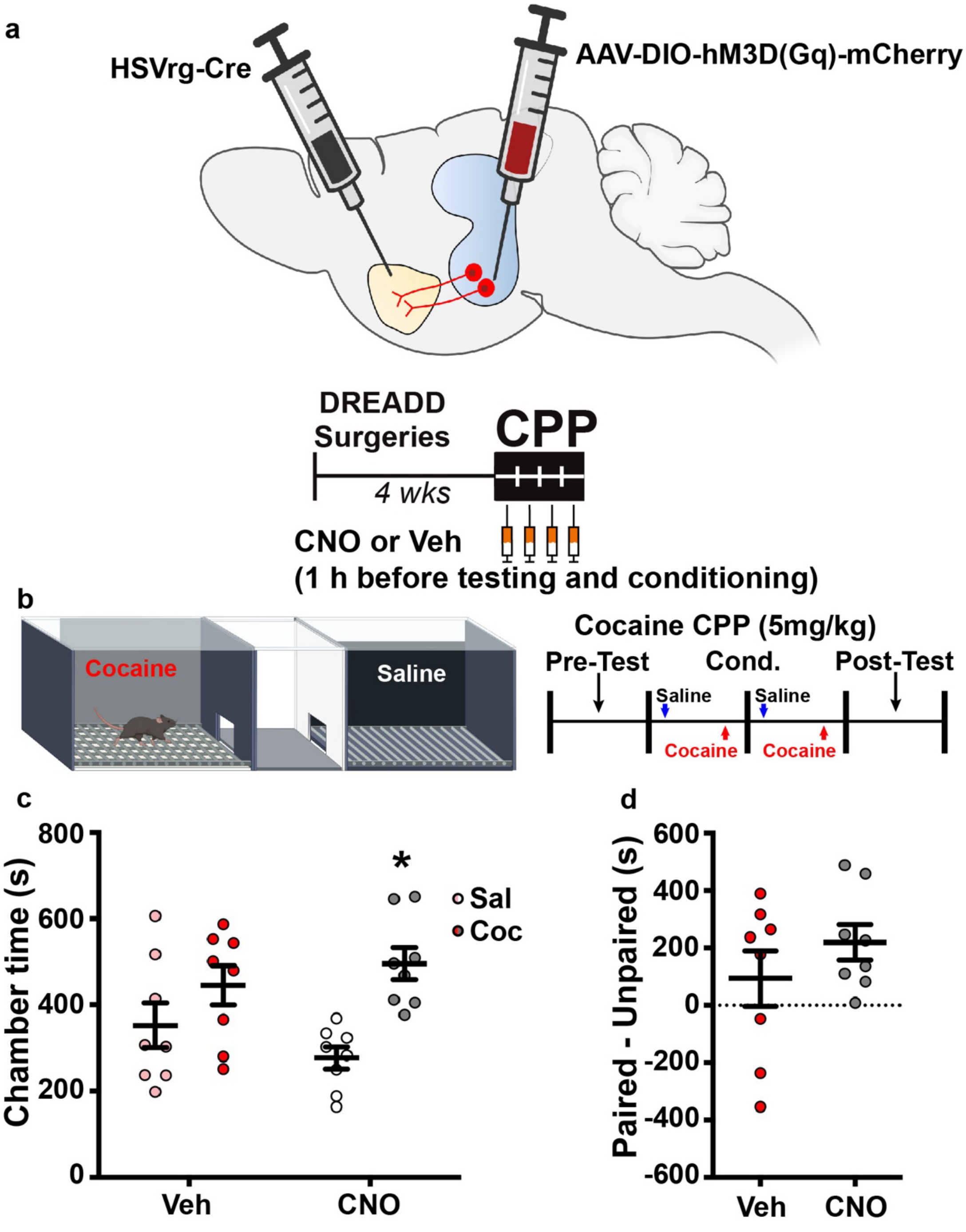

**Supplemental Figure 5.**

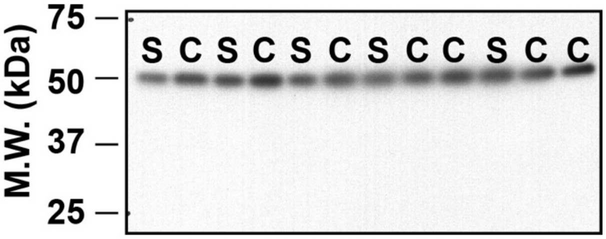
